# Characterization and contribution of RPE senescence to Age-related macular degeneration in *Tnfrsf10* knock out mice

**DOI:** 10.1101/2023.08.04.552052

**Authors:** Iori Wada, Kenichiro Mori, Parameswaran G Sreekumar, Rui Ji, Christine Spee, Elise Hong, Keijiro Ishikawa, Koh-Hei Sonoda, Ram Kannan

**Affiliations:** Doheny Eye Institute, Pasadena, CA, USA; Department of Ophthalmology, Graduate School of Medical Sciences, Kyushu University, Fukuoka 812-8582, Japan; Stein Eye Institute, Geffen School of Medicine, University of California, Los Angeles, CA, USA

**Keywords:** TNFRS10A mutation, RPE, senescence, aging, mitochondrial function, PKC, senolytics

## Abstract

**Background:** Retinal pigment epithelial cells (RPE) play vital role in the pathogenesis of age-related macular degeneration (AMD). Our laboratory has shown that RPE cellular senescence contributed to the pathophysiology of experimental AMD, and SASP members are involved in this process. Recently, we presented confirmatory evidence to earlier GWAS studies that dysregulation of tumor necrosis factor receptor superfamily 10A (TNFRSF10A) dysregulation leads to AMD development and is linked to RPE dysfunction. This study aims to investigate the contribution of RPE senescence to AMD pathophysiology using *TNFRSF10A* silenced human RPE (hRPE) cells and Tnfrsf10 KO mice.

**Methods:** Sub-confluent primary hRPE cells and *TNFRSF10A* silenced hRPE were exposed to stress-induced premature senescence with H2O2 (500 μM, 48h), and senescence-associated markers (βgal, p16, and p21) were analyzed by RT-PCR and WB analysis. The effect of H2O2-induced senescence in non-silenced and silenced hRPE on OXPHOS and glycolysis was determined using Seahorse XF96 analyzer. Male C57BL/6J Tnfrsf10 KO (*Tnfrsf10^-/-^*) mice were used to study the regulation of senescence by TNFRSF10A *in vivo*. Expression of p16 and p21 in control and KO mice of varying ages were determined by RT-PCR, WB, and immunostaining analysis.

**Results:** The senescence-associated p16 and p21 showed a significant (*p* < 0.01) upregulation with H2O2 induction at the gene (1.8- and 3-fold) and protein (3.2- and 4-fold) levels in hRPE cells. The protein expression of p16 and p21 was further significantly increased by co-treatment with siRNA (*p* < 0.05 vs. H2O2). Mitochondrial oxygen consumption rate (OCR) and extracellular acidification rate (ECAR) (pmol/min/total DNA) increased with senescence induction by H2O2 for 48h in control RPE, and knockdown of *TNFRSF10A* caused a further increase in OCR and ECAR. In addition, co-treatment with PKC activator significantly improved all parameters. Similarly, *in vivo* studies showed upregulation of p16 and p21 by RT-PCR, WB, and immunostaining analysis in RPE/choroid of Tnfrsf10 KO mice. When subjected to examination across distinct age groups, namely young (1-3 months), middle (6-9 months), and old (12-15 months) mice, a discernible age-related elevation in the expression of p16 and p21 was observed.

**Conclusions:** Our findings suggest that TNRSF10A is a regulator of regulates in RPE senescence. Further work on elucidating pathways of senescence will facilitate the development of new therapeutic targets for AMD.

## 1. Introduction

Age-related macular degeneration (AMD), the predominant cause of irreversible visual impairment in the elderly population, is characterized by the degeneration of the retinal pigment epithelium (RPE) and photoreceptors [1, 2]. Advanced age, environmental influences, and genetic predisposition exert considerable influence over the likelihood of developing AMD. Extensive genome-wide association studies (GWAS) have identified more than 34 susceptibility loci associated with AMD [3]. We have previously reported the notable involvement of mutations in *TNFRSF10A* as a preeminent genetic predisposition for the pathological condition [4, 5].

In our previous investigation, we demonstrated the expression of TNFRSF10A in retinal pigment epithelium (RPE) cells within the human ocular framework [6]. Employing siRNA-mediated knockdown of *TNFRSF10A* in human RPE (hRPE) cells, we observed a decline in cellular viability and an upsurge in apoptotic events, attributed to the concurrent downregulation of protein kinase C-α (PKCA) expression [6]. This evidence supports the notion that diminished TNFRSF10A expression triggers RPE degeneration, thereby potentially contributing to the pathogenesis of AMD. Notably, our findings also substantiated the development of photoreceptor dysfunction and RPE degeneration, reminiscent of key clinical manifestations of AMD, in *Tnfrsf10*^-/-^ mice. Given the recent findings of the emerging role of senescence in RPE dysfunction and AMD development [7, 8], it was of interest to delineate the association between TNFRSF10A and cellular senescence and investigate the underlying mechanisms.

Aging is widely acknowledged as a crucial predisposing factor in the pathogenesis of AMD. The process of aging leads to diverse structural and physiological alterations in the human retina [9]. Numerous independent investigations have indicated that aging contributes to the onset of various ocular disorders [10, 11]. One such study discovered a substantial loss of RPE cells in the peripheral region of the human retina [12]. Furthermore, this study demonstrated that the proportion of RPE to photoreceptor cells diminishes throughout the retina with advancing age [13]. Moreover, protein levels of senescence markers p16, p21, and p53, were elevated in RPE cells derived from aged human donors [14]. Long-term *in vitro* cultures of RPE cells also exhibit signs of senescence [15], and the presence of SA-β-galactosidase, an indicator of cellular senescence, has been reported in RPE cells of both human and primate retinas [16]. Intracellular deposition of lipofuscin in RPE cells can also trigger the generation of reactive oxygen species (ROS) upon exposure to oxygen and light [17]. We hypothesize that the senescence of RPE cells plays a crucial role in the pathogenesis of age-related retinal diseases and potential of targeting these cells could become an effective therapeutic strategy for AMD.

The aim of this study is to investigate the involvement of TNFRSF10A in the process of cellular senescence, thereby proposing a therapeutic modality for AMD that specifically targets TNFRSF10A-mediated cellular senescence.

## 1. MATERIALS AND METHODS

### 2.1 Retinal pigment epithelial cell culture

All procedures were conducted in accordance with the principles outlined in the Declaration of Helsinki pertaining to Research Involving Human Subjects. Retinal pigment epithelial (RPE) cells were extracted from human fetal ocular tissues obtained from Advanced Bioscience Resources Inc. (Alameda, CA, USA) and Novogenix Laboratories, LLC (Los Angeles, CA, USA). The cells were then cultured as previously described [8]. In brief, human RPE (hRPE) cells were cultured in Dulbecco’s modified Eagle’s medium (DMEM, #15-013-CV, Corning, NY, USA) supplemented with 10% fetal bovine serum (FBS, #4800-500HI, Laguna Scientific, Laguna Niguel CA, USA). Subculturing of cells was performed, and passages 2 to 4 were utilized for all experiments once confluency was attained.

### 2.2 Transfection

The human retinal pigment epithelial (hRPE) cells were cultivated in 6- or 96-well plates, devoid of any antibiotics reaching a cellular confluence of 30-50% over a period of 24 hours.. For transfection purposes, either a scrambled control siRNA (4390843, Invitrogen) or TNFRSF10A siRNA (NM_003844 SASI_Hs01_00139577, Merck, Darmstadt, Germany) was combined with a specialized RNA transfection reagent (Lipofectamine RNAi MAX; 13778075, Invitrogen) and Opti-MEM (31985-070, Grand Island, NY, USA), following the manufacturer’s protocol. These mixtures were then diluted using Dulbecco’s modified Eagle’s medium (DMEM; D6046, Sigma-Aldrich), supplemented with 1% fetal bovine serum (FBS) and 1% penicillin/streptomycin (P4333, Sigma-Aldrich). Following a 24-hour period post-transfection, the composite transfection mixture was carefully replaced with DMEM in preparation for subsequent assays.

### 2.3 Induction of cellular senescence

Senescence was induced in RPE using two stimuli as described below.

#### 2.3.1. H2O2-induced RPE senescence

Sub confluent human RPE cells were treated with 500 μM H2O2 (Sigma-Aldrich #H1009, Saint Louis, MO, USA) after sci control or siRNA transfection. The H2O2 treatment was repeated the following day. The medium was replaced, cells were washed in PBS and incubated with fresh medium containing 10% FBS. Cells were kept for another 48 h, and medium was replaced every 24 h.

#### 2.3.2. Doxorubicin (DOX) induced senescence

Subconfluent (50–60% confluent at the initiation of the study), human RPE cells were subjected to treatment with 0.2 μg/ml doxorubicin (DOX) (Sigma-Aldrich, #D1515, St. Louis, MO, USA) for a duration of 3 hours in a culture medium containing 0.5% fetal bovine serum (FBS). Subsequently, the cells were washed with PBS and transitioned to fresh medium supplemented with 10% FBS.

### 2.4 Senescence-associated β-galactosidase (SA-β gal) assay

The SA-βgal activity was assessed in RPE cells at the end of the experiment using the Senescence Detection Kit (Sigma-Aldrich, #CS0030) according to the manufacturer’s instructions. Briefly, cells were seeded in 4-well chamber slides, washed with PBS at the end of the experiment, and fixed with 4% PFA for 10 min at room temperature. Cells were washed with PBS and incubated overnight at 37℃ without CO2 and with β-gal substrate. After washing, cells were examined with a microscope and photographed (Keyence, Itasca, IL, USA). The expression of blue dye around the nucleus was taken as an indicator of senescent cells. For quantification, blue-stained cells and total cells were counted under a microscope, and the percentage of cells expressing β-galactosidase (senescent cells) was calculated [18].

### 2.5 RNA Isolation and Real-Time Quantitative RT-PCR

Total RNA was extracted utilizing an RNA isolation kit (Qiagen), and reverse transcription polymerase chain reaction (RT-PCR) was executed following established protocols [19]. The expression levels of genes were standardized against *tubulin* mRNA and expressed as fold-change compared to the control. The primer sequences employed in the experiment can be found in Table 1. The relative fold-change in mRNA expression was determined by computing 2-^ΔΔCT^. The outcomes are depicted as the average disparity in relative fold-change in mRNA expression ± standard error of the mean (SEM).

### 2.6 Western Blot Analysis

Protein extraction was performed from cells and the posterior eyecup, excluding conjunctiva, sclera, and muscle tissue. RIPA buffer supplemented with a protease inhibitor was employed for this purpose. The concentration of soluble protein was determined using BSA as a standard (SpectraMax iD5, Molecular Devices, Sunnyvale, CA, USA). Equal quantities of protein (30 - 60 μg) were resolved on TGX-precast gels (Bio-Rad Laboratories Inc., Hercules, CA, USA) and subsequently transferred to PVDF blotting membranes (Millipore, Billerica, MA, USA). The membranes were incubated with respective primary antibodies overnight at 4 C (The antibodies and dilutions used are listed in Table 2). Following incubation with suitable secondary antibodies (Vector Laboratories, Burlingame, CA, USA), protein bands were visualized using a chemiluminescence (ECL) detection system (Bio-Rad Laboratories Inc., Hercules, CA, USA). Adequate protein loading was confirmed by probing for GAPDH.

### 2.7 Measurement of Cellular Respiration

Mitochondrial bioenergetics was determined by measuring oxygen consumption rate (OCR) of RPE cells with a Seahorse XFe96 Analyzer (Agilent, Santa Clara, CA, USA [18]. Sub-confluent hRPE cells were induced senescence with H2O2 (500 μM) induction with sci control or siRNA transfection. Assays were initiated by replacing the growth medium with a 175 μl XF assay medium. XF assay medium contained glucose (25 mM), sodium pyruvate (1 mM), and glutamine (2 mM). The concentration of inhibitors was oligomycin (ATP-Synthase inhibitor) at 1.5μM; carbonyl cyanide 4-(trifluoromethoxy) phenylhydrazone (FCCP, mitochondrial membrane depolarizer) at 1.5 μM, and a mixture of 0.5 μM of each rotenone (complex 1 inhibitor) and antimycin A (complex 3 inhibitors). We calculated basal respiration, maximal respiration, OCR-linked ATP production, spare respiratory capacity, non-mitochondrial oxygen consumption, and proton leak. The experiments were repeated three times. The OCR data were expressed as pmol/min/µg protein.

### 2.8 Measurement of Cellular Glycolysis

The Seahorse XF Glycolysis Stress Test Kit (Agilent, #103020-100) was utilized following the standard protocol established by Agilent Seahorses, as previously described [8]. This kit enables the assessment of glycolytic pathway capacity subsequent to glucose deprivation. During glycolysis, the resulting acidification of the surrounding cell medium is directly measured by the analyzer and reported as the Extracellular Acidification Rate (ECAR). Human retinal pigment epithelial cells were seeded in 96-well plates and induced into senescence following treatment with H2O2 (500 μM), with separate groups subjected to sci control or siRNA transfection. To investigate the glycolytic potential of hRPE cells after the treatment period, the ECAR was measured in 17-20 wells per experimental condition before and after sequential injections of glucose (10 mM), oligomycin (1 μM), and 2-Deoxy-D-glucose (2-DG; 50 mM). Data were expressed as milli-pH per minute per microgram of protein (mpH/min/µg protein).

### 2.9 Detection of Mitochondrial Superoxide with MitoSOX

Superoxide production within the mitochondria was visualized using MitoSOX (Thermo Fisher Scientific, Waltham, MA, USA). hRPE cells cultured on four-well chamber slides at subconfluent densities were subjected to treatment with *TNFRSF10A* siRNA or / and H2O2 alone or co-administered with PKC activator (PMA). Prior to the termination of the treatment, cells were incubated with 5 μM MitoSOX for 15 minutes at 37 ◦C. Subsequently, the cells were rinsed with PBS, fixed with 3.7% PFA for 15 minutes, washed again, mounted using 4′,6-diamidino-2phenylindole (DAPI: Vector Laboratories, Burlingame, CA, USA), and examined employing a laser confocal microscope (LSM 710: Zeiss, Thornwood, NY, USA).

### 2.10 Animals

All animal experiments were conducted in strict accordance with the guidelines set forth by the Association for Research in Vision and Ophthalmology (ARVO) Statement for the Utilization of Animals in Ophthalmic and Visual Research. Wild type C57BL/6 J mice (procured from Charles River Laboratories Japan, Yokohama, Japan) and Tnfrsf10^-/-^ mice at the respective ages of 3 weeks or 12 months were used in our studies. The Tnfrsf10^-/-^ mice (RRID: MMRRC_030532-MU) were procured from the Mutant Mouse Resource and Research Center (MMRRC) situated at the University of Missouri, which is an NIH-supported repository for genetic strains. These specific mice strains were generously contributed to the MMRRC by Wafik S. El-Deiry, M.D., Ph.D., affiliated with the University of Pennsylvania. To ensure genetic stability, the Tnfrsf10-/-mice utilized in our study were selectively bred with C57BL/6 J mice for a minimum of five generations within our laboratory. The strains employed in our studies were meticulously screened through DNA sequencing to confirm the absence of the rd8 mutation.

### 2.11 TEM

The posterior segments of murine eyes were subjected to fixation in a solution containing 4% glutaraldehyde and 0.1 M cacodylate buffer at a temperature of 4°C, undergoing overnight incubation. Following rinsing with 0.1 M cacodylate buffer, the samples were subsequently post-fixed for a duration of 1.5 hours, utilizing a mixture comprising 1% OsO4 and 0.1 M cacodylate buffer. After this step, dehydration was accomplished through an ethanol series, ultimately leading to the embedding of the samples within an epoxy resin. The polymerization process took place at 45°C for a duration of 30 minutes, followed by an extended period of 72 hours at 60°C. To facilitate the analysis of retinal structure, semi-thin sections measuring 1 µm were meticulously sliced from the resin blocks employing a Reichert Ultracut S microtome (Leica Microsystems Inc., Buffalo Grove, IL, USA). These sections were subsequently stained with Azure II. By employing a light microscope, a suitable region of interest within the sample was carefully identified for further examination using transmission electron microscopy (TEM). For TEM imaging, ultra-thin sections with thicknesses ranging from 60 to 70 nm were produced using the Reichert Ultracut S microtome. These sections were mounted onto copper grids and subjected to staining with a solution composed of 0.2% oolong tea extract and an EM stainer (Nissin EM, Tokyo, Japan) before being subjected to observation utilizing a TEM instrument (HT7700; Hitachi, Tokyo, Japan).

### 2.12 Immunofluorescence

Paraffin sections (5 μm thick) were fixed with methanol for 20 minutes and washed with PBS. After a 30-minute blocking step with 10% goat serum, the tissues were incubated overnight at 4℃ with primary antibodies, p21 (sc-6246), p16^INK4A^ (sc-56330), and then incubated with corresponding secondary antibodies (Vector Laboratories Inc., Newark, CA, USA). Images were acquired using Keyence (BZ-X710, KEYENCE, Osaka, Japan). The quantification of the images was according to a previous report [20].

### 2.13 Statistical Analysis

All statistical analysis were performed using graphing software (Prism 8; GraphPad Software). All data are expressed as mean ± SEM. Statistical significance was determined by unpaired two-tailed Student’s *t* test or Analysis of variance (ANOVA), followed by Dunnett’s multiple comparison test. *P* values < 0.05 were considered significant.

## 3 Results

### 3.1 Upregulation of β-gal positive senescent cells with H2O2 or DOX and TNFRSF10A knockdown

To evaluate the hypothesis that attenuation of TNFRSF10A engenders the onset of cellular senescence, we utilized a well-established model wherein subconfluent hRPE cells were subjected to a 500 μM H2O2 treatment as described in our previous study [18]. Oxidative stress induced by H2O2 with this protocol yielded a marked escalation in cellular senescence (*p* < 0.01 vs. controls). Furthermore, the concurrent administration of *TNFRSF10A* siRNA and H2O2 elicited a significant, further augmentation of cellular senescence; conversely, treatment with siRNA alone failed to show a statistically significant increase in comparison to control group. Moreover, the concomitant administration of a PMA at a concentration of 100 nM effectively abrogated the progression of cell senescence. In addition to the aforementioned model, we also conducted studies using the anthracycline antibiotic doxorubicin (DOX), another model used for inducing cellular senescence [8]. Consistent with the findings of the DOX study, co-treatment with TNFRSF10A siRNA led to a substantial increase in the population of SA-β-gal positive cells. Furthermore, incubation in the presence of PMA resulted in a marked reduction in the number of SA-β-gal positive cells.

### 3.2 Increased expression of senescence-related biomarkers with H2O2 and upregulation with TNFRSF10A knockdown

Next, we assessed senescence-associated genes and proteins in RPE senescence using RT-PCR and WB analysis. Treatment with H2O2 along with *TNFRSF10A* siRNA resulted in a significant upregulation in the expression of senescence markers, cyclin-dependent kinase inhibitor 1 (p21) and -2A (p16) mRNA, in comparison to the control group. Moreover, co-treatment with *TNFRSF10A* siRNA and H2O2 caused a substantial elevation in p21 and p16 mRNA expression levels, which was higher than H2O2 treatment alone group. Correspondingly, the expression levels of p16 and p21 proteins were significantly increased in senescent cells subjected to H2O2 treatment, and the knockdown of *TNFRSF10A* further elevated their levels by 5-8 fold when compared to the control group. Notably, the co-treatment of H2O2 and *TNFRSF10A* siRNA in the presence of PMA significantly suppressed the expression of p21 and p16 at both mRNA and protein levels. In summary, our findings suggest that *TNFRSF10A* siRNA alone does not induce appreciable senescence. However, it effectively promotes senescence in RPE cells that are already undergoing senescence and inhibits senescence by facilitating PKC signaling.

### 3.2 Mitochondrial biogenesis and energetics are altered by H2O2-induced senescence

Mitochondrial fission is orchestrated by DRP1 [21], which relocates from the cytosol to the outer mitochondrial membrane, where it engages with receptor proteins, including fission protein-1 (Fis1) [22]. In this study, we aimed to explore whether H2O2-induced cellular senescence upholds mitochondrial dynamics through the regulation of DRP1. Senescent cells exhibited a significant upregulation in DRP1 protein expression. Moreover, the co-treatment of *TNFRSF10A* siRNA resulted in a more pronounced elevation in protein levels, an effect that was counteracted by co-treatment with PMA (Figure 3). A comparable pattern was observed for Fis1, where its expression decreased upon PMA treatment. Notably, senescent cells displayed an increasing trend in Mfn2 expression, suggesting alterations in mitochondrial fusion.

**Figure 1.**
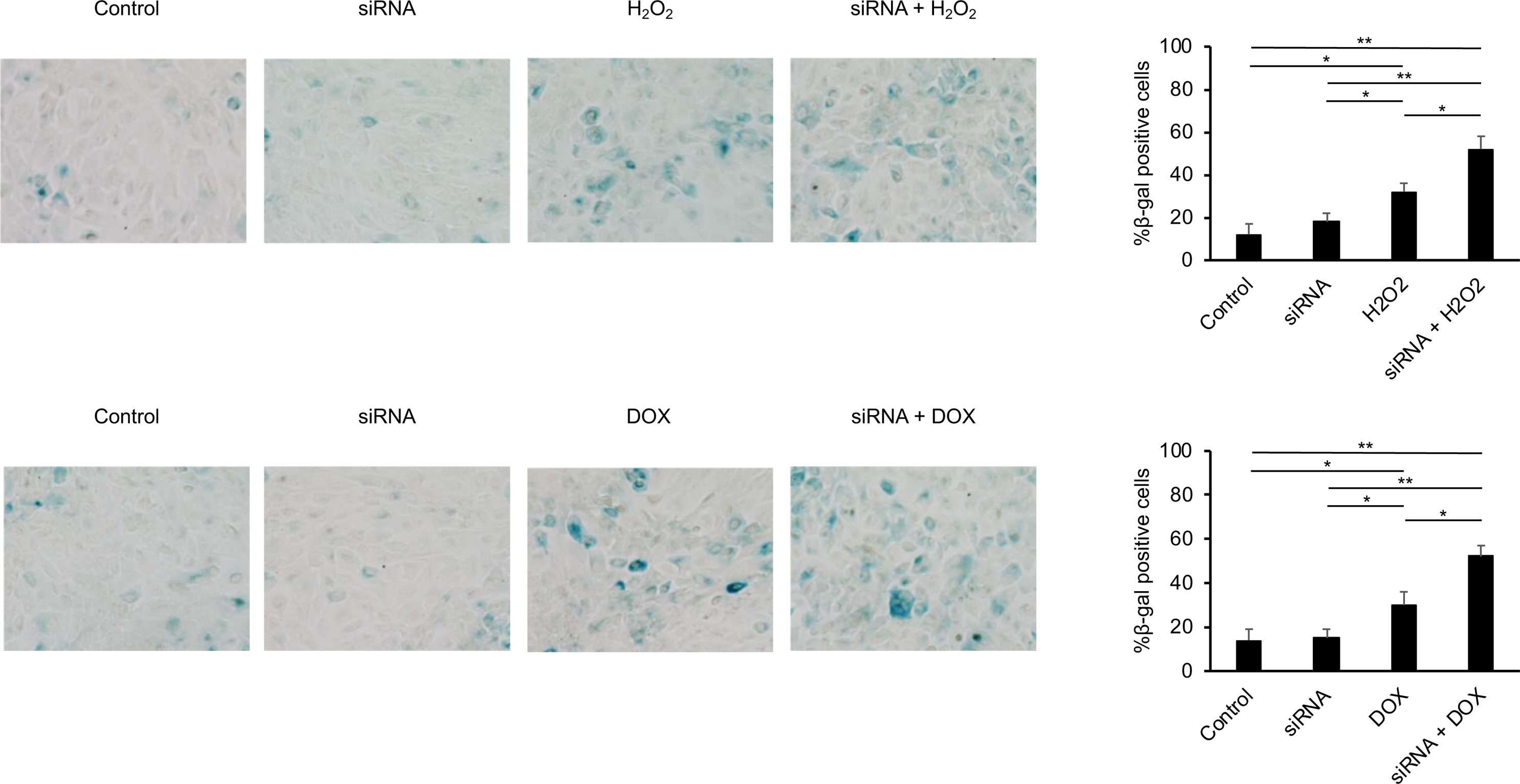
Upregulation of H2O2 and doxorubicin (DOX)-induced cellular senescence (β-galactosidase positive cells) by *TNFRSF10A* siRNA. Co-treatment with *TNFRSF10A* siRNA significantly augmented both H2O2-induced senescence (Figures A and B) and DOX-induced senescence (Figures C and D). RPE cells were treated with H2O2 (500 μM) in a serum-free medium for 2 hours, and the medium was subsequently replaced with a 10% FBS-containing medium. Sub-confluent hRPE cells were also exposed to DOX (0.2 μg/ml) for 3 hours in a medium containing 0.5% FBS. Subsequently, the medium was then changed to a 10% FBS medium for 2 days. Samples were processed for β-galactosidase staining. The oxidative stress significantly augmented the number of β-galactosidase-positive cells, and the senescence of RPE cells was notably enhanced by the pre-administration of *TNFRSF10A* siRNA (Figures B and D). The scale bar represents 50 μm. \**p* < 0.05 and \*\**p* < 0.01. n = 3.

**Figure 2.**
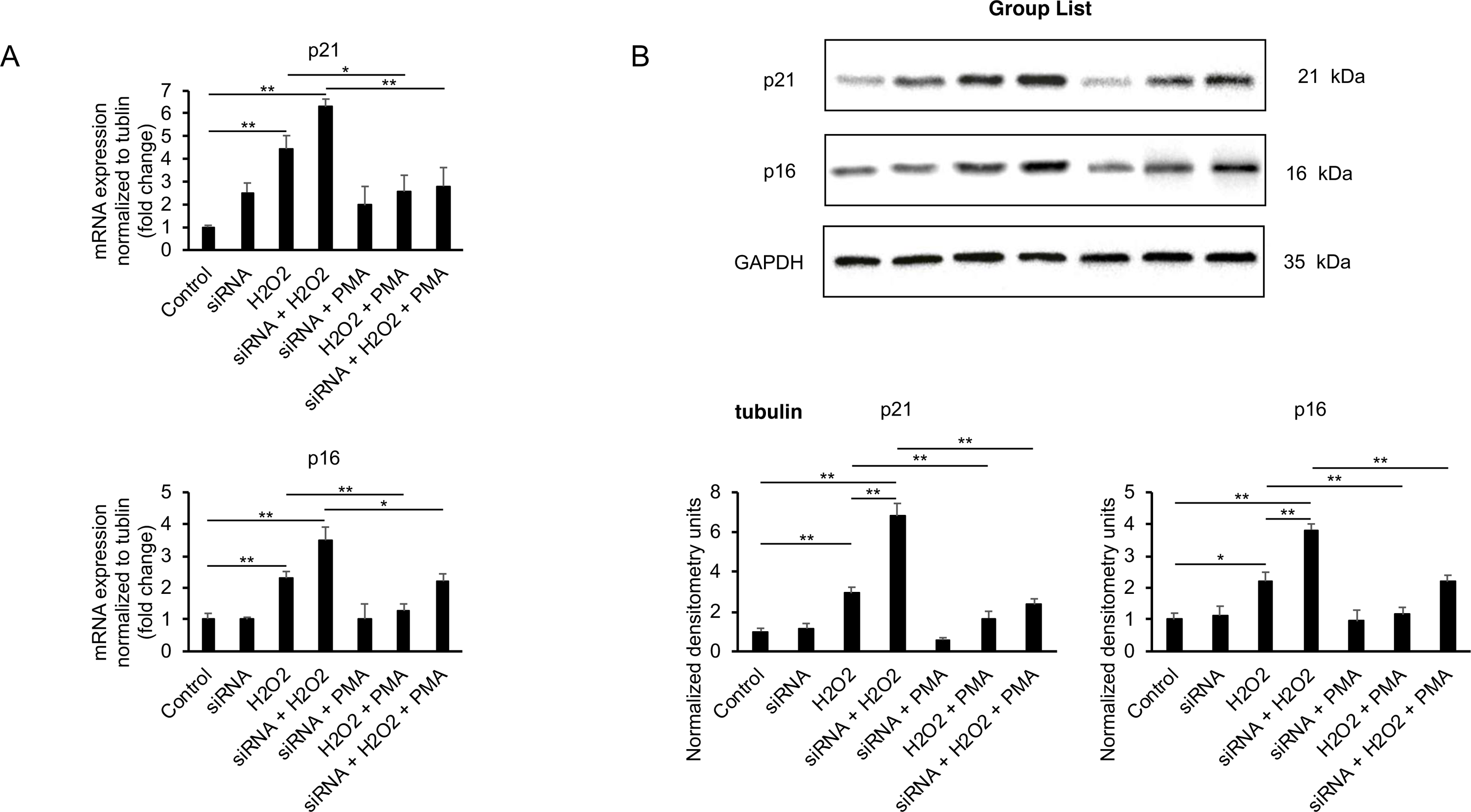
Suppression of senescence in H2O2-induced senescence through the activation of protein kinase C (PKC). The expression of p21 and p16 at both mRNA (A) and protein (B) levels exhibited a notable increase during the process of senescence. *TNFRSF10* siRNA further accelerated the senescence phenotype. However, treatment with a PKC activator significantly attenuated the aging process. Western blots were subjected to densitometry analysis and subsequently normalized to GAPDH expression. \**p* < 0.05 and \*\**p* < 0.01. n = 3-4.

**Figure 3.**
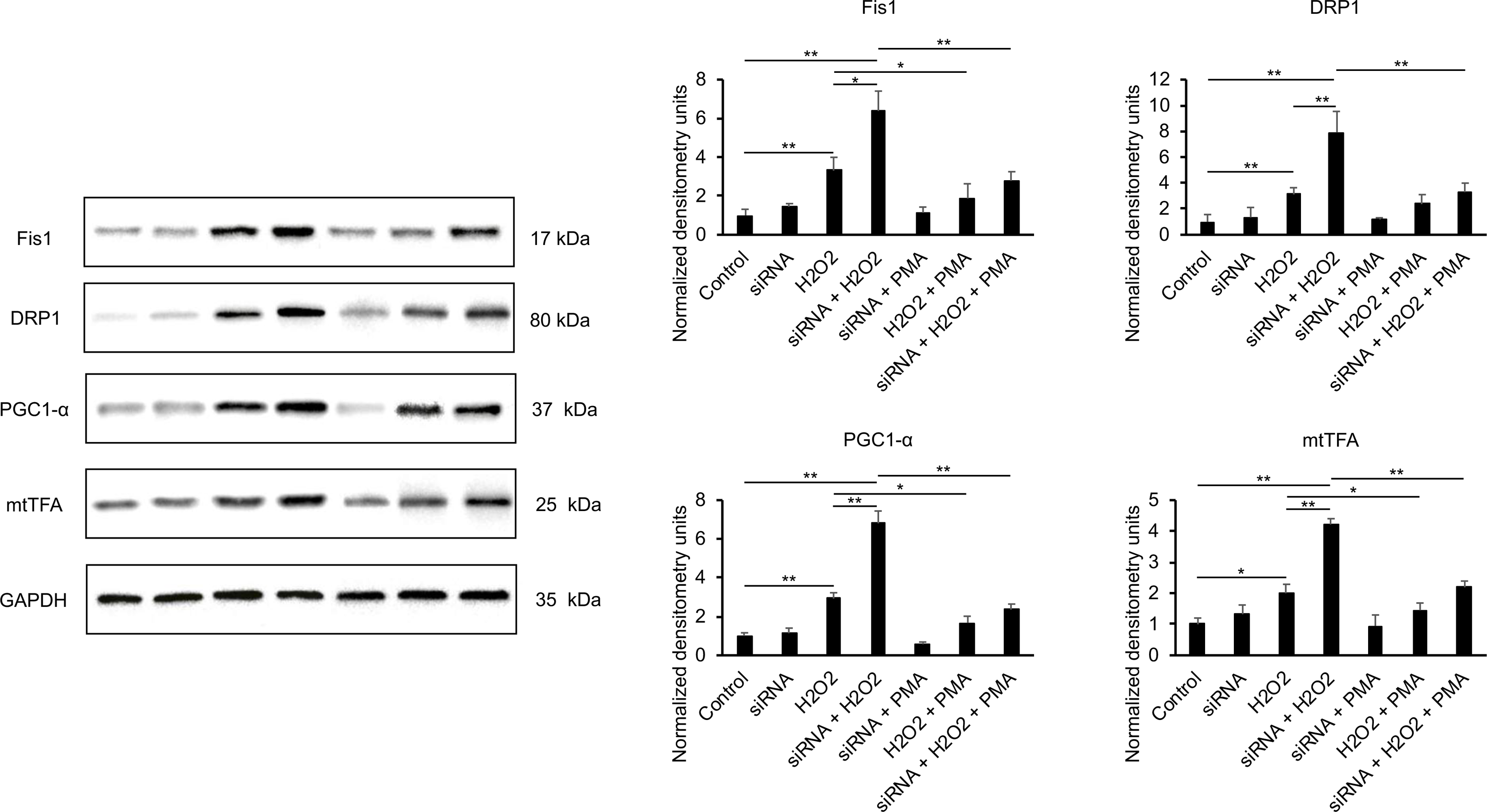
Regulation of mitochondrial biogenesis and fission-fusion proteins by a PKC activator in senescent hRPE cells. Sub-confluent RPE cells were subjected to treatment with 500 μM H2O2 alone or in combination with TNFRSF10A siRNA. Protein expression levels were assessed using WB analysis. The bar graphs presented here represent the quantitative densitometry analysis derived from three independent experiments, displayed as mean ± standard deviation (SD). Senescence was accompanied by a significant increase in the expression of mitochondrial fission proteins, namely Fis1 and DRP1. However, treatment with the PKC activator markedly inhibited the expression of these proteins and mitigated the senescent phenotype. Notably, RPE senescence was associated with a significant elevation in the levels of biogenesis markers, such as PGC1 and mtTFA, whereas co-treatment with PKC activators led to a decrease in their expression. Conversely, no substantial changes were observed in the levels of the mitochondrial fusion protein Mfn2. \**p* < 0.05 and \*\**p* < 0.01. n = 3.

Furthermore, we investigated the involvement of PGC-1α, a known participant in the transcriptional regulation of mitochondrial biogenesis and respiratory function [23]. Additionally, the mitochondrial transcription factor A (mtTFA) emerged as another crucial factor governing mitochondrial biogenesis, playing a role in mtDNA transcription initiation and the packaging of mtDNA into nucleoids [24]. We observed a significant increase in the protein expression of PGC1α, as well as a 1.8-fold elevation in mtTFA levels in H2O2-induced senescent cells. Furthermore, co-treatment with siRNA led to a significant augmentation in their respective levels.

Taken together, these findings suggest that senescence in RPE cells may coincide with an activation of mitochondrial biogenesis, and the influence of PMA may modulate these processes.

### 3.3. Modulation of Energy Metabolism and the Impact of PKC Activator in Senescent RPE Cells

Senescence is often accompanied by alterations in energy metabolism, which can vary depending on the cell type and inducer [25–28]. In our study, we investigated mitochondrial function in H2O2-induced senescence. Senescent cells exhibited enhanced ATP production, elevated maximal respiration, and increased spare respiratory capacity when compared to non-senescent cells (Figure 4). Furthermore, co-treatment with *TNFRSF10A siRNA* significantly augmented basal respiration compared to non-aging cells, as well as maximal respiration, ATP production, and spare respiratory capacity when compared to aging cells. These findings indicate that senescent cells exhibit heightened mitochondrial ATP synthesis compared to their non-senescent counterparts. Notably, co-treatment of RPE cells with a PKC activator reduced all mitochondrial bioenergetic parameters to the level observed in non-senescent cells (Figure 4).

**Figure 4.**
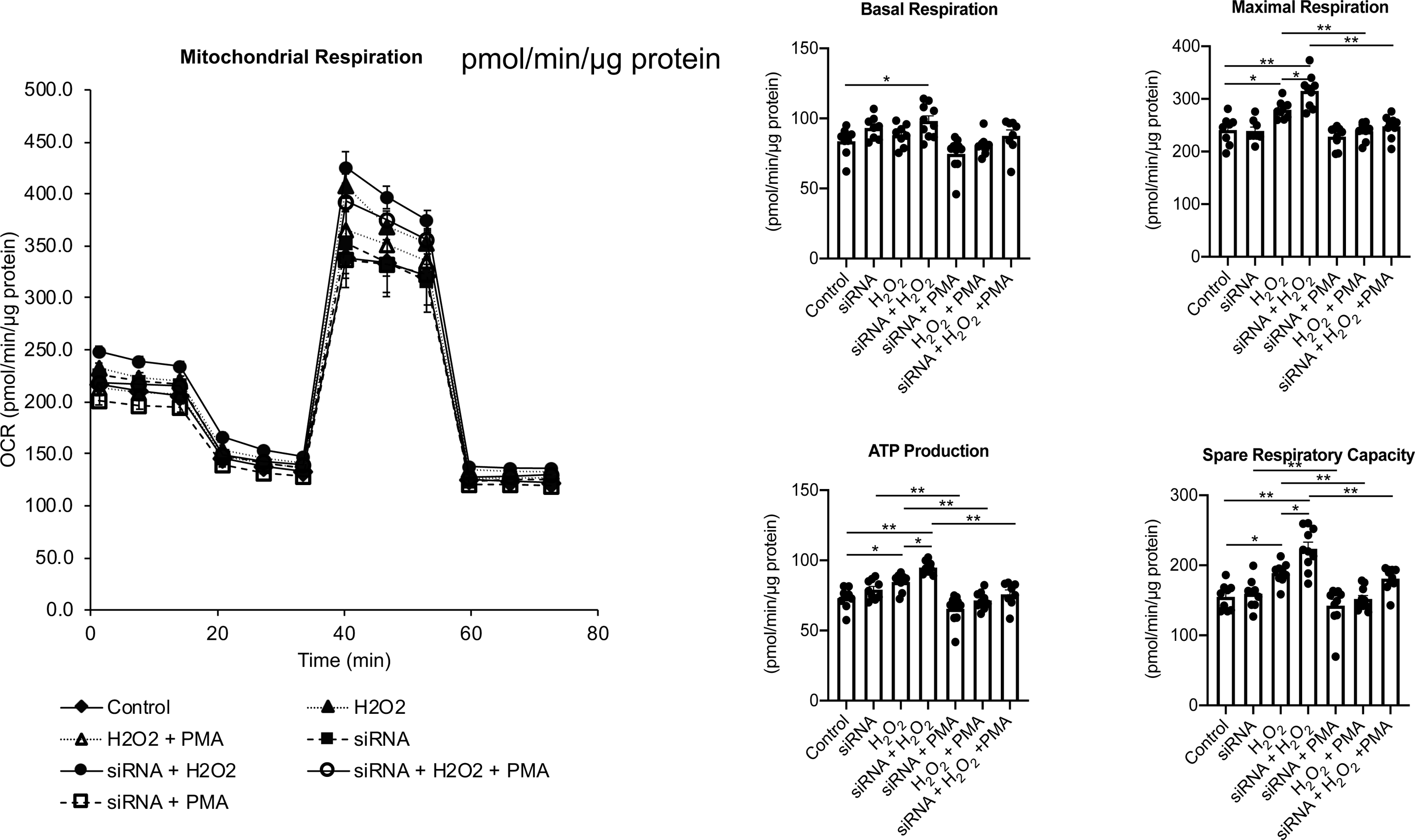
Enhanced OCR parameters during senescence and their attenuation by a PKC activator. Mitochondrial bioenergetics were evaluated using Seahorse XFe96 analysis. Senescence induced by H2O2 in hRPE cells resulted in a significant increase in maximal respiration, ATP production, and spare respiratory capacity. Moreover, co-treatment with *TNFRSF10A* siRNA further augmented these parameters. However, the concurrent administration of PKC activators exhibited a significant decrease in these senescent cells. Values are presented as means ± standard error (SE) (n=8-10). \**p* < 0.05 and \*\**p* < 0.01. n = 8-12.

Considering the substantial increase in mitochondrial function observed during RPE senescence, it is reasonable to expect alterations in the main catabolic pathways. Therefore, we investigated changes in glycolysis by assessing the extracellular acidification rate (ECAR) under conditions of glucose starvation followed by subsequent addition. Senescent cells displayed elevated ECAR, increased glycolytic capacity, and enhanced glycolytic reserve compared to control cells (Figure 5). Furthermore, co-treatment with *TNFRSF10A* siRNA significantly heightened all these parameters. These findings suggest that senescent cells exhibit an augmented contribution to glycolysis compared to non-senescent cells. To examine whether senescence induces changes in glycolysis, we employed a PKC activator to inhibit senescence progression. Co-treatment with the PKC activator substantially reduced glycolytic function in senescent cells. Hence, senescent RPE cells rely relatively on both glycolysis and oxidative phosphorylation (OXPHOS).

**Figure 5.**
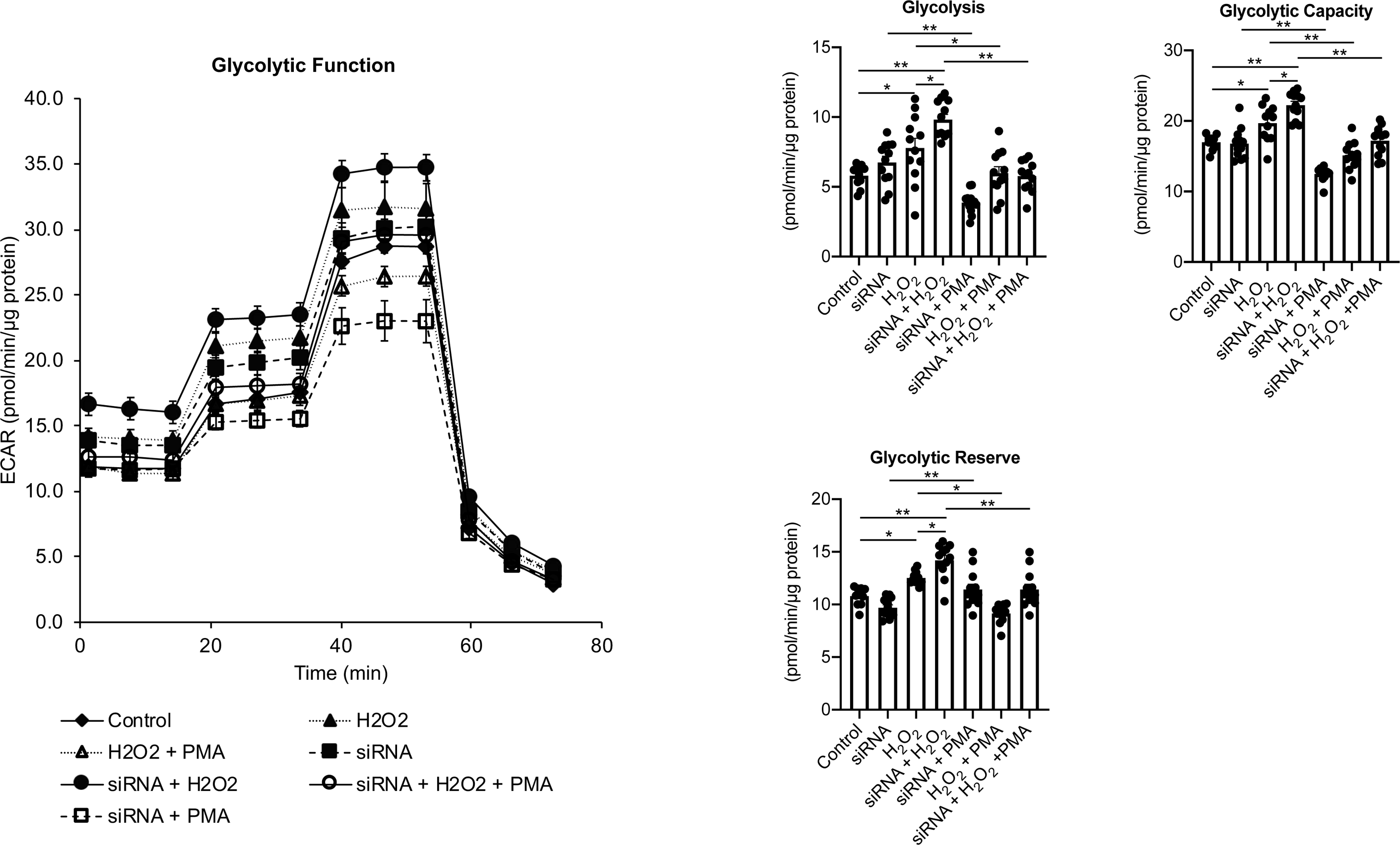
Alterations in glycolysis in H2O2-induced senescent hRPE cells. The glycolytic function in senescent-induced hRPE cells was assessed by real-time monitoring using the Seahorse XFe96 Glycolytic Stress Test Kit, which measures key parameters of glycolysis. H2O2-induced senescence in hRPE cells resulted in a significant increase in glycolysis, glycolytic capacity, and glycolytic reserve. Notably, co-treatment with *TNFRSF10A* siRNA further augmented these glycolytic parameters. However, the administration of PKC activators exerted a significant suppressive effect on all glycolytic functions. The data were normalized by total cellular protein, and the values are presented as means ± standard error of the mean (SEM). \**p* < 0.05 and \*\**p* < 0.01. n = 9 – 15

### 3.4. Inhibition of mitochondrial ROS production in senescent-hRPE cells exposed to TNFRSF10A siRNA by PKC activator treatment

ROS are widely implicated in mitochondrial DNA damage and are believed to play a crucial role in the pathogenesis of AMD [29]. Thus, we sought to investigate whether H2O2 induces oxidative stress in hRPE cells under our conditions of H2O2-induced senescence. We observed that H2O2 exposure resulted in the generation of mitochondrial superoxide in senescent hRPE cells. Furthermore, co-treatment with *TNFRSF10A* siRNA significantly augmented mitochondrial superoxide production. Conversely, administration of the PKC activator effectively suppressed ROS generation induced by H2O2, both in the presence and absence of *TNFRSF10A* siRNA (Figure 6).

**Figure 6.**
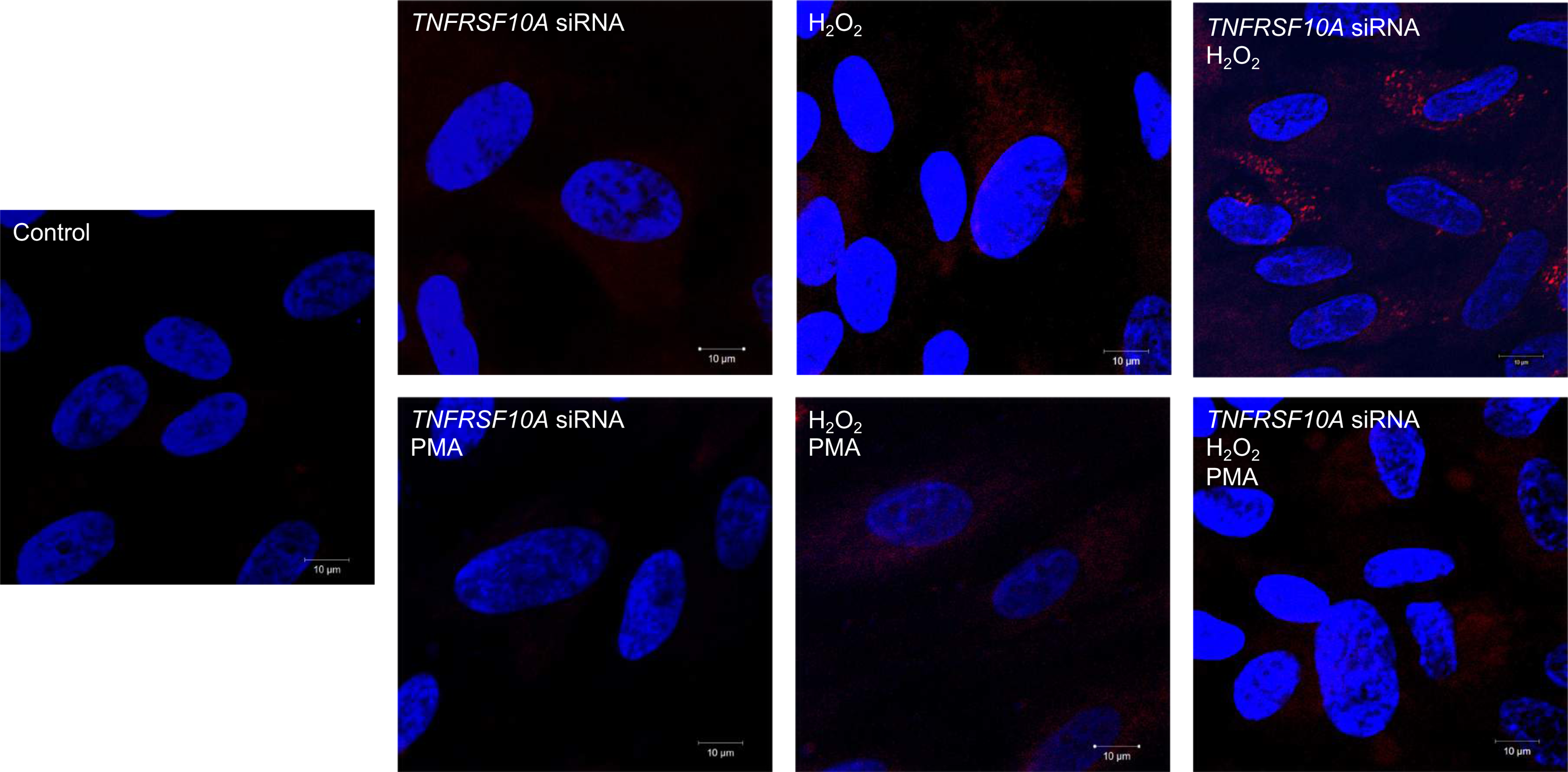
Mitochondrial ROS (mtROS) production in H2O2- and/or *TNFRSF10A* siRNA-induced senescent hRPE cells. To examine mtROS production in senescent hRPE cells induced by 500 μM H2O2 and/or *TNFRSF10A* siRNA, the cells were cultured in 4-well chamber slides. After the respective treatments, the cells were incubated with 5µM MitoSox Red, a marker for mitochondrial superoxide, for 10 minutes at 37℃. Following a washing step, the cells were imaged using ZEISS LSM 710 microscopy. As illustrated in the figure, a significant increase in mtROS levels was observed in H2O2-treated cells co-treated with TNFRSF10A siRNA. Moreover, co-treatment with PKC activators significantly decreased ROS production. Red: Mitochondrial superoxide, Blue, DAPI, nuclear stain. *Scale bar*: 10 µm.

### 3.5. TNFRSF10A siRNA triggers the activation of SASP-related genes in hydrogen peroxide H2O2-induced senescence of RPE cells

To elucidate the activation of SASP-related genes during senescence in hRPE cells, we subjected the cells to treatment with H2O2 alone or in combination with *TNFRSF10A* siRNA, followed by analysis through RT-PCR. Senescent cells display metabolic activity and manifest SASPs, which encompass a diverse array of inflammatory cytokines, chemokines, metalloproteinases, and growth factors [30–32]. These factors are known to exert paracrine effects, thereby modulating and governing tissue homeostasis. The SASP has garnered attention as both a potential instigator and a therapeutic target for various age-related conditions, such as AMD [33]. Thus, we investigated the SASP within the context of our experimental parameters.

Conditioned medium (CM) was collected from both control and senescent cells. Exposure to H2O2 significantly elevated the expression of the inflammatory mediator-related gene *IFN-ɣ* and *IL1-β* (*p* < 0.01, compared to non-aging control cells). Notably, co-treatment with *TNFRSF10A* siRNA substantially augmented the expression of these mediators (*p* < 0.01) (Figure 7A). Among the interleukins, *IL-6* gene were prominently expressed in senescent cells relative to non-aging cells (Figure 7B). Furthermore, co-treatment with *TNFRSF10A* siRNA significantly upregulated the expression of all interleukins compared to the control group (*p* < 0.01). It is well known that matrix metalloproteinases (MMPs) play a pivotal role in the regulation of extracellular matrix (ECM) turnover and maintenance of structural integrity [34]. Co-treatment with *TNFRSF10A* siRNA led to a marked increase in the expression of both *MMP1* and *MMP3* (*p* < 0.01, Figure 7C).

**Figure 7.**
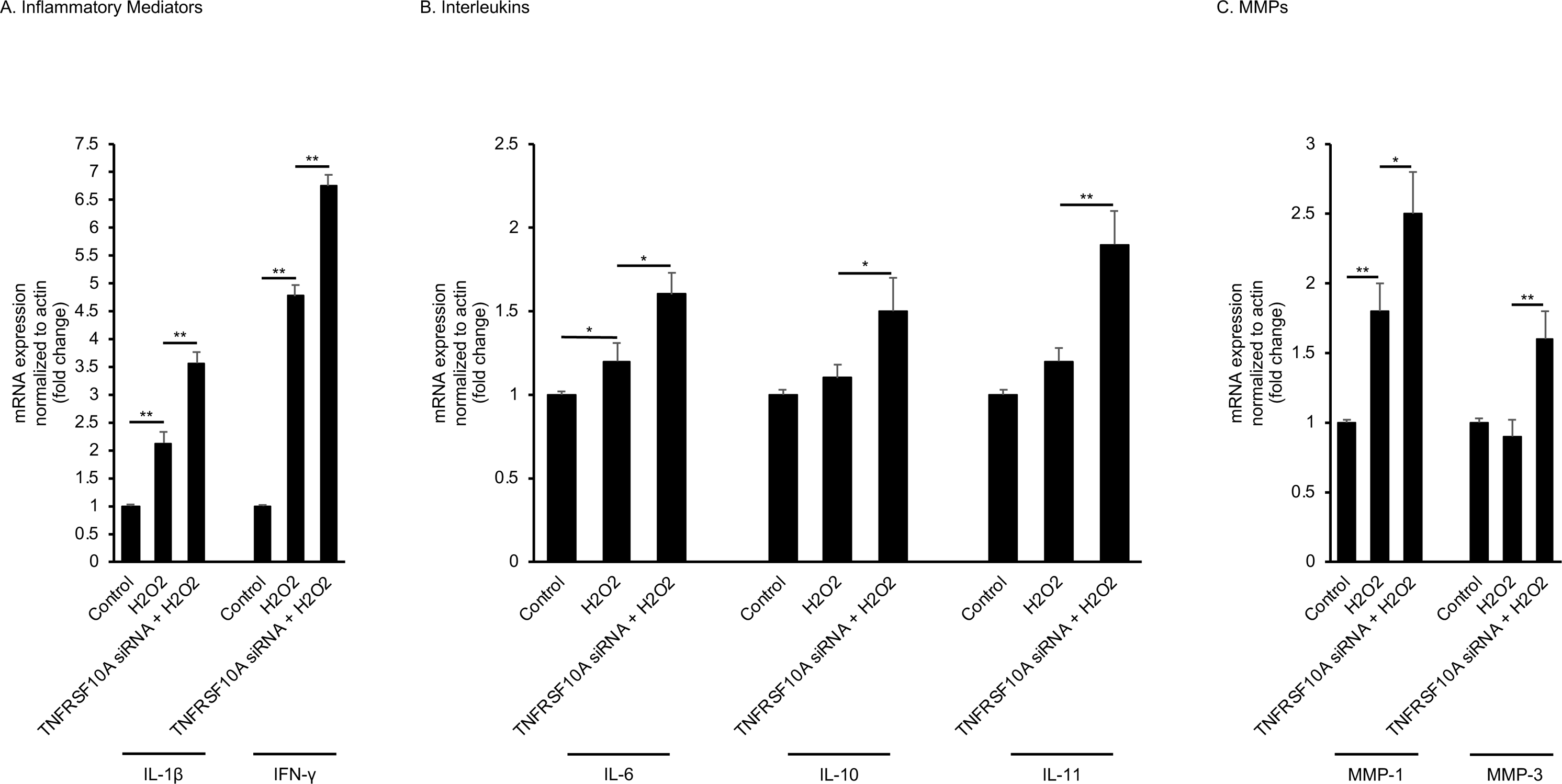
Effect of TNFRSF10A siRNA on the secretion of SASP members in senescent RPE cells. Subconfluent human RPE cells were treated with 500 μM H2O2 alone or with TNFRSF10A siRNA. The H2O2 treatment was repeated the following day. The RPE-conditioned medium was collected at 2 h and assayed by RT-PCR. Data are converted to fold change relative to controls and are derived from three samples / experimental conditions. TIMPs-tissue inhibitors of metalloproteinases, MMPs-Matrix metalloproteinases. \**p* < 0.05 and \*\**p* < 0.01. n = 4

### 3.6. Mitochondrial Abnormalities in Tnfrsf10^-/-^ Mice

To investigate the association between aging and senescence mediated by *Tnfrsf10*, we employed *Tnfrsf10^-/-^* mice, which were previously characterized by our research group [6]. Our analysis focused on assessing mitochondrial ultrastructural changes using transmission electron microscopy (TEM). At 6 months of age, *Tnfrsf10^-/-^*mice exhibited a combination of hypertrophied mitochondria suggestive of hyperfusion and fragmented mitochondria indicative of fission when compared to wild-type (*WT*) mice (Figure 8). These findings indicate a dysregulation of mitochondrial dynamics in the absence of *Tnfrsf10*. Moreover, upon reaching 12-months, *WT* mice mitochondria displayed a structure nearly identical to that observed at 6 months. In contrast, *Tnfrsf10*^-/-^ mice at 12 months exhibited notable mitochondrial hyperfusion and hypertrophy, indicating a potential involvement of Tnfrsf10 in mitochondrial dysfunction with aging. Further systematic studies will be required to understand the correlation between aging and senescence (see also below).

**Figure 8.**
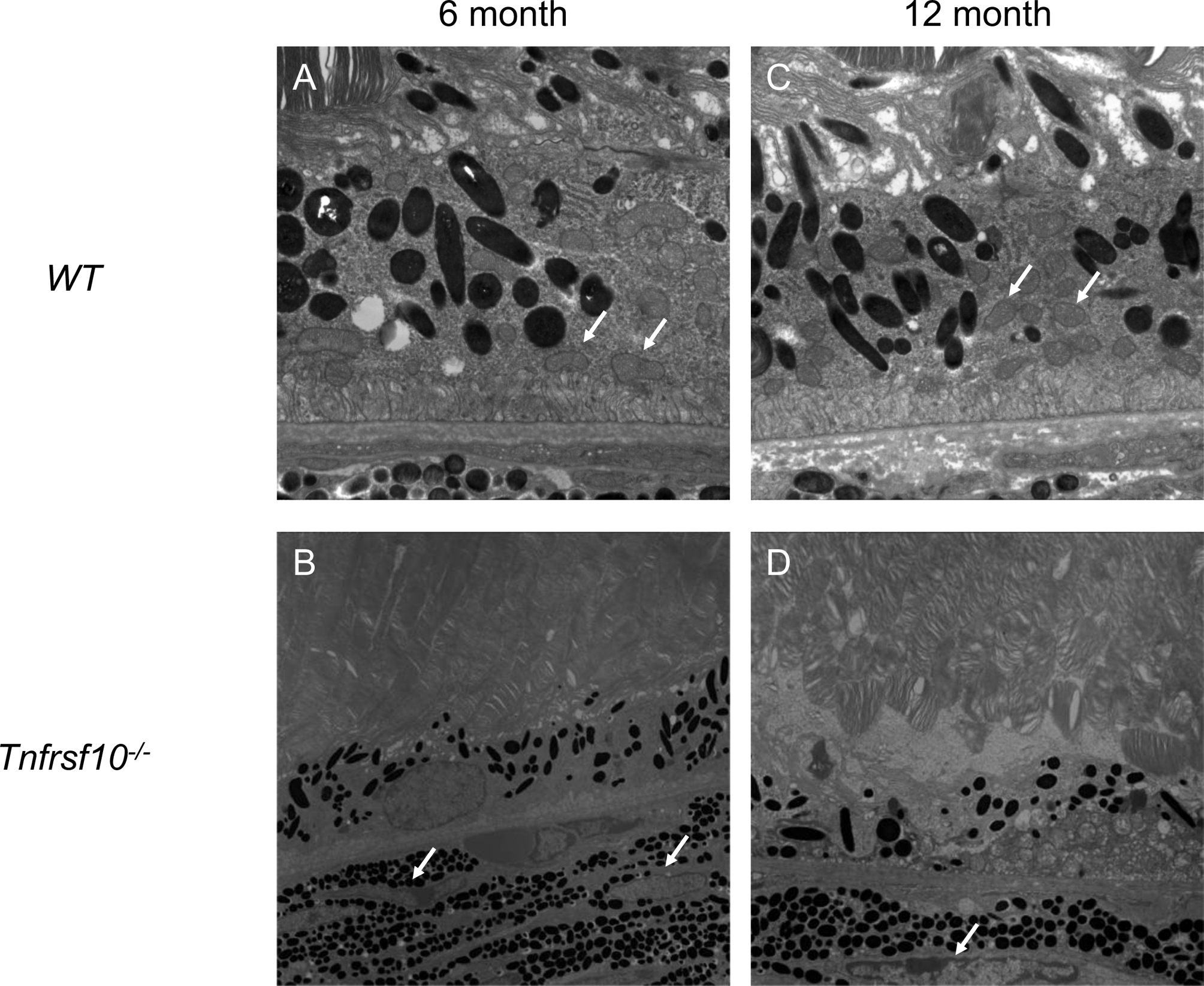
Representative images of transmission electron microscopy of the RPE from 6M and 12 M *WT* (A, C) or *Tnfrsf10^-/-^* mice (B, D). White arrowheads indicate the mitochondria.

### 3.7. Elevation of Aging-Related Factors in Tnfrsf10^-/-^ Mice

In this study, we investigated the effect of alterations in RPE layer and mitochondrial dysfunction on the aging process in 12-month-old *WT* and *Tnfrsf10^-/-^* mice. p21 and p16 were expressed substantially higher in the RPE layer and choroid of *Tnfrsf10*^-/-^ mice compared to their *WT* mice (Figure 9A). Similarly, our investigations using WB analysis with RPE and choroidal cells showed these findings (Figure 9B). These results indicated that our evidence points to apotential involvement of *Tnfrsf10* in modulating the expression of senescence factors through its influence on mitochondrial function.

**Figure 9.**
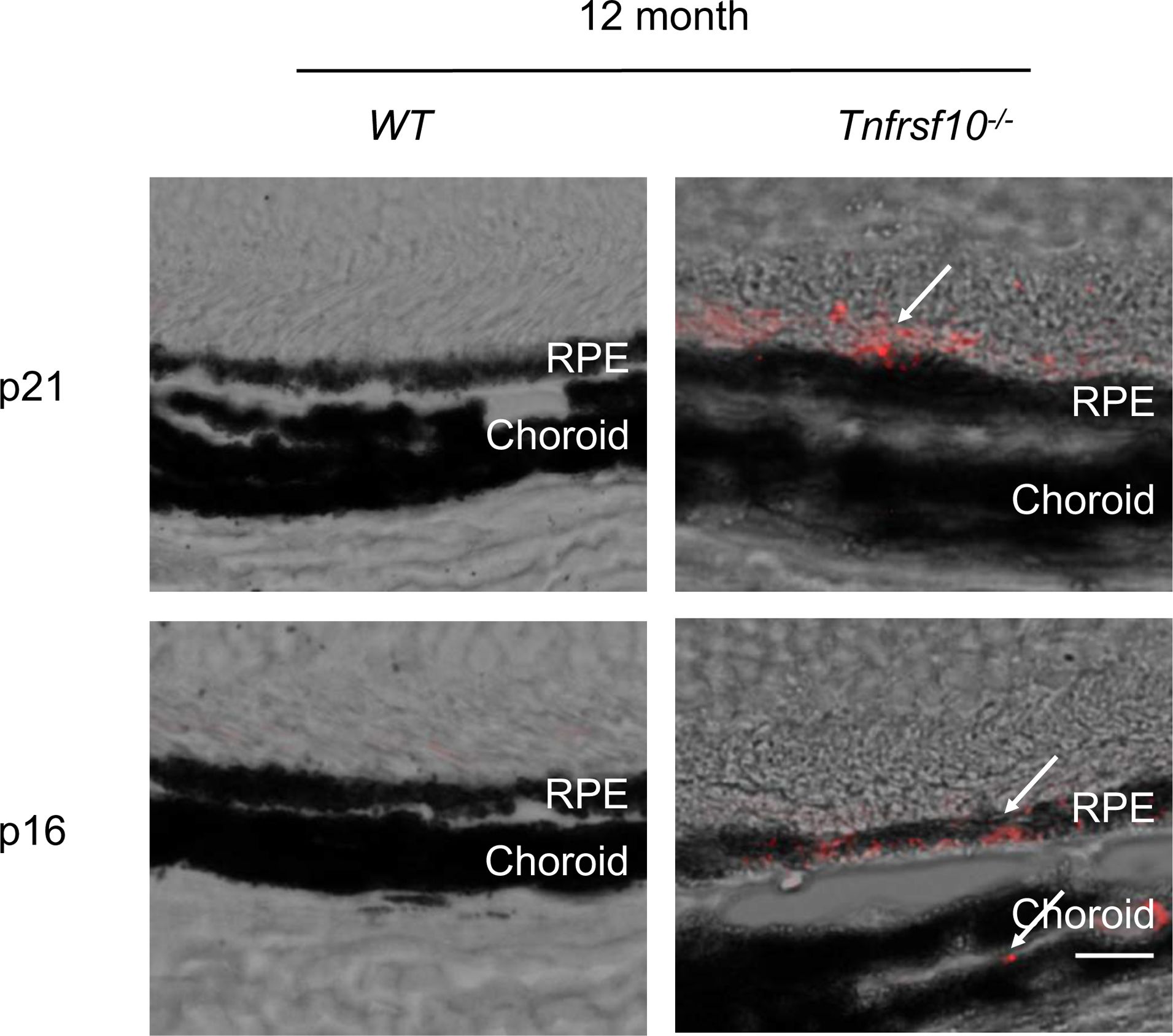
Evidence for senescence in RPE cells in *Tnfrsf10A*^-/-^ mice. Evidence for the presence of senescent RPE cells by p21 and p16 expression in RPE and choroid from 12M old mice. White arrows show positive immunoreactivity for p21 and p16. RPE-Retinal pigment epithelium.

### 3.8. Age-associated upregulation of senescence-related factors

To investigate the alterations in the expression of senescence-related factors within the RPE layer during the aging process, we examined *Tnfrsf10^-/-^* mice at distinct stages of life: young (1-3 months), middle-aged (6-9 months), and old age (12-15 months). Intriguingly, in young mice, the expression of senescence-related factors, namely p21 and p16, was detected solely in the neuroretinal layer, while their expression was negligible or below detection in the RPE layer. However, in middle-aged and old mice, both p21 and p16 became prominently expressed within the RPE layer, with a notable accumulation of expression observed in the old mice (Figure 10A). Moreover, upon investigating aging-associated factors by WB utilizing RPE and choroidal tissue as our focus, a striking age-dependent upregulation in protein expression was unveiled (Figure 10B). These results suggest that the expression of aging-related factors in the RPE layer is affected by aging in *Tnfrsf10^-/-^* mice.

**Figure 10.**
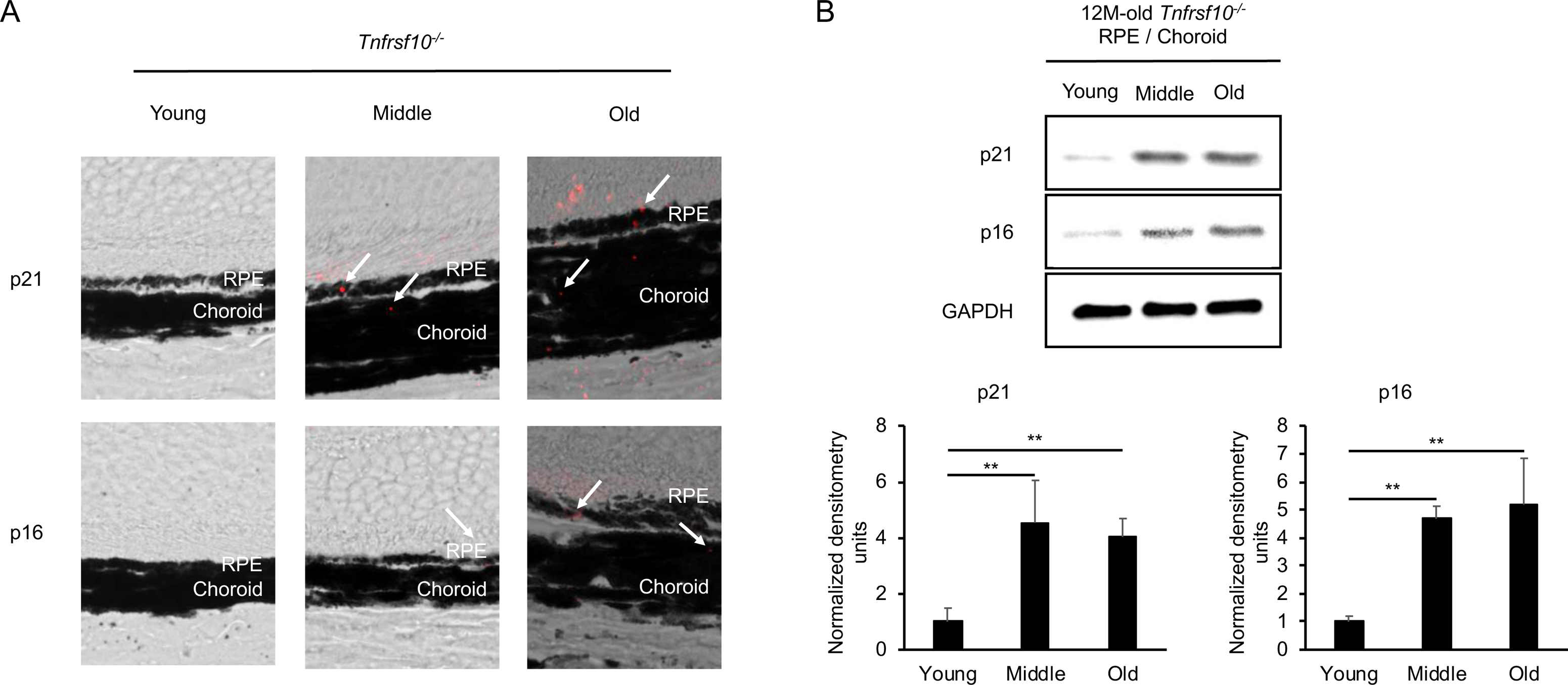
The relationship between senescence in RPE cells and aging. Senecent RPE cells by p21 and p16 expression in RPE and choroid from aged mice (A). White arrows show positive immunoreactivity for p21 and p16. RPE-Retinal pigment epithelium.The expression of p21 and p16 at protein levels exhibited a notable increase during the aging (B). Western blots were subjected to densitometry analysis and subsequently normalized to GAPDH expression. \**p* < 0.05 and \*\**p <* 0.01. n = 3

**Figure 11.**
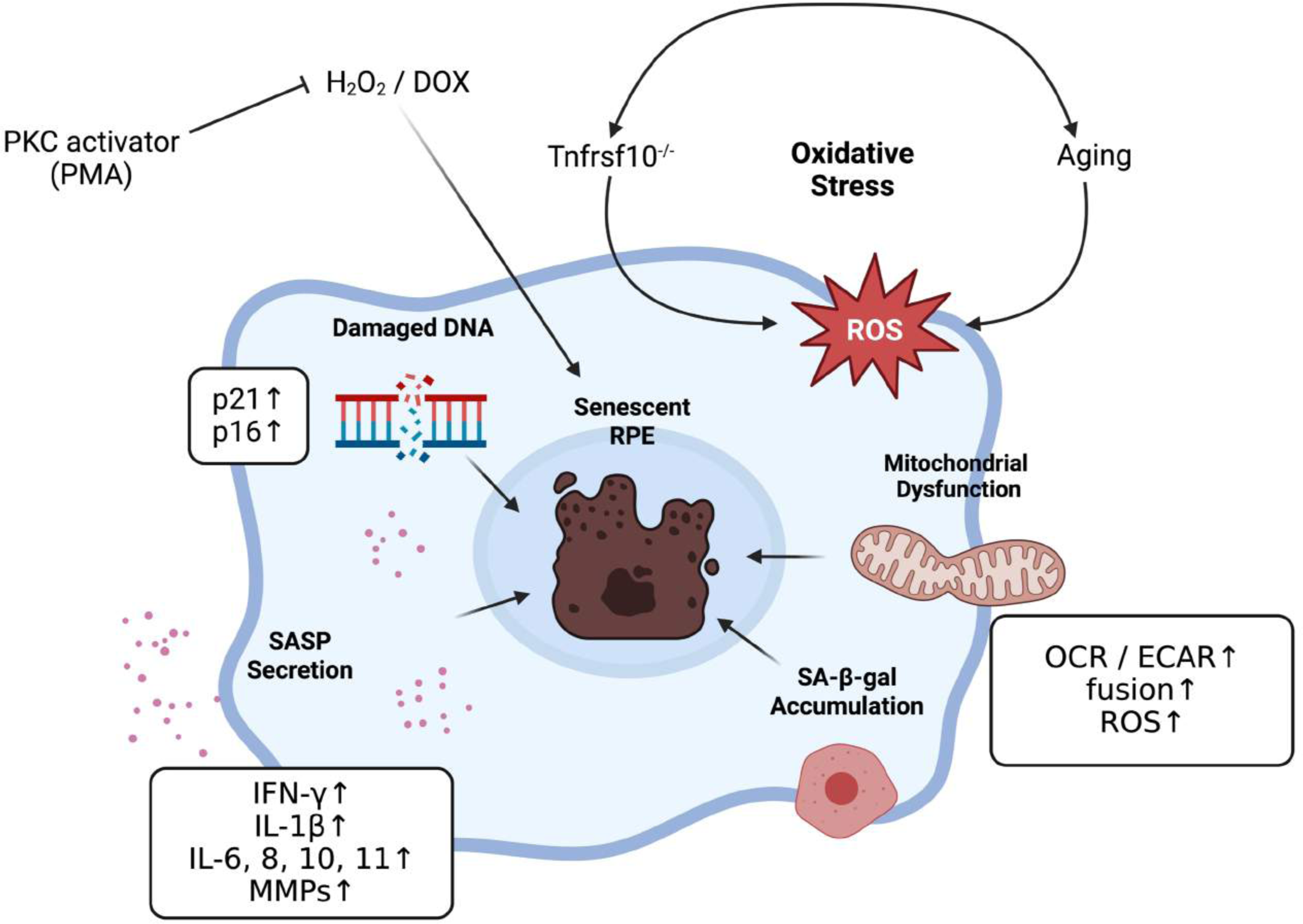
The scheme of this study.

## 4. Discussion

Environmental and genetic risk factors play a pivotal role in exacerbating oxidative stress, thereby contributing to the progression of AMD [35–39]. Notably, aging emerges as a significant risk factor for AMD, instigating the accumulation of oxidized proteins, lipids, and DNA [40, 41]. Moreover, compelling evidence suggests a noteworthy association between the presence of high-risk alleles for complement factor H (CFH), a major AMD susceptibility factor, and a substantially heightened level of mitochondrial DNA (mtDNA) damage in affected donors [42].

In this study, our focus was on *TNFRSF10A*, a genetic factor implicated in AMD. We conducted investigations on senescence-related factors in hRPE cells in vitro, employing *TNFRSF10A* siRNA and H2O2. Intriguingly, although treatment with *TNFRSF10A* siRNA alone did not yield any notable differences compared to the control group, when co-treated with H2O2, the induction of cellular senescence was significantly augmented. Our findings also were confirmed by in vivo studies utilizing *Tnfrsf10^-/-^* mice, as the mice advanced in age, a notable augmentation in the expression p16 and p21 was observed at the RPE layer.

Previous studies have highlighted the importance of oxidative stress and damage as crucial triggers for RPE degeneration. Oxidative stress primarily stems from an excess production of ROS within the mitochondria [35, 43]. In line with this, our study revealed an increase in ROS production from the mitochondria in the group subjected to co-treatment with *TNFRSF10A* siRNA and H2O2. Additionally, we observed an upregulation of PGC1-α expression in the group subjected to co-treatment with *TNFRSF10A* siRNA and H2O2. PGC1-α is known to promote the expression of antioxidants that offer protection against ROS-induced mitochondrial damage and cell death [44]. Furthermore, PGC1-α governs the expression of numerous genes crucial for mitochondrial biogenesis, which plays a pivotal role in energy production [38, 39]. The observed elevation in OCR and ECAR under *TNFRSF10A* siRNA and H2O2 stimulation in our study may represent a cellular response to the excessive stress induced by mitochondrial dysfunction. This suggests that the increase in OCR and ECAR could be a compensatory mechanism initiated in response to the heightened stress associated with mitochondrial dysfunction.

This study provides evidence of the role of TNFRSF10A in promoting mitochondrial dysfunction and age-related cellular senescence. AMD is a chronic inflammatory disease characterized by persistent oxidative stress impacting RPE cells [45]. In normal conditions, the ubiquitin-proteasome and lysosomal/autophagy pathways play vital roles in eliminating damaged proteins and organelles [46]. However, in RPE with AMD, proteostasis is compromised due to the accumulation of lysosomal lipofuscin and extracellular drusen [47]. In previous study, the PKC activator employed demonstrated significance in activating the ubiquitin-proteasome system [48], which may explain its senolytic properties in this study. However, further investigations are necessary as PKC activators also impact the cell cycle. This is because cellular senescence is the arrest of a stable cell cycle, which in normal cells is triggered in response to various intrinsic and extrinsic stimuli and developmental signals [49, 50]. Continued research will shed light on these mechanisms and their potential therapeutic applications in AMD.

## 5. Conclusions

The current findings suggest that *TNFRSF10A* is intricately involved in the cellular senescence process, exerting its influence through the regulation of mitochondrial function in RPE. Moreover, these results clearly demonstrate the accelerated progression of cellular senescence in the context of aging.

## Supporting information

Table for primer

## Acknowledgments

We thank Ernesto Barron and Eric Barron for their help in using the KEYENCE BZ-X-700 Imaging System.

This work was supported by NIH/NEI grants R01EY30141, a gift from Keck Foundation to DEI.

This work was presented at the Annual ARVO meeting (IOVS April 2023, Vol.64, 5384.)

---

**Supplemental Figure 1.**
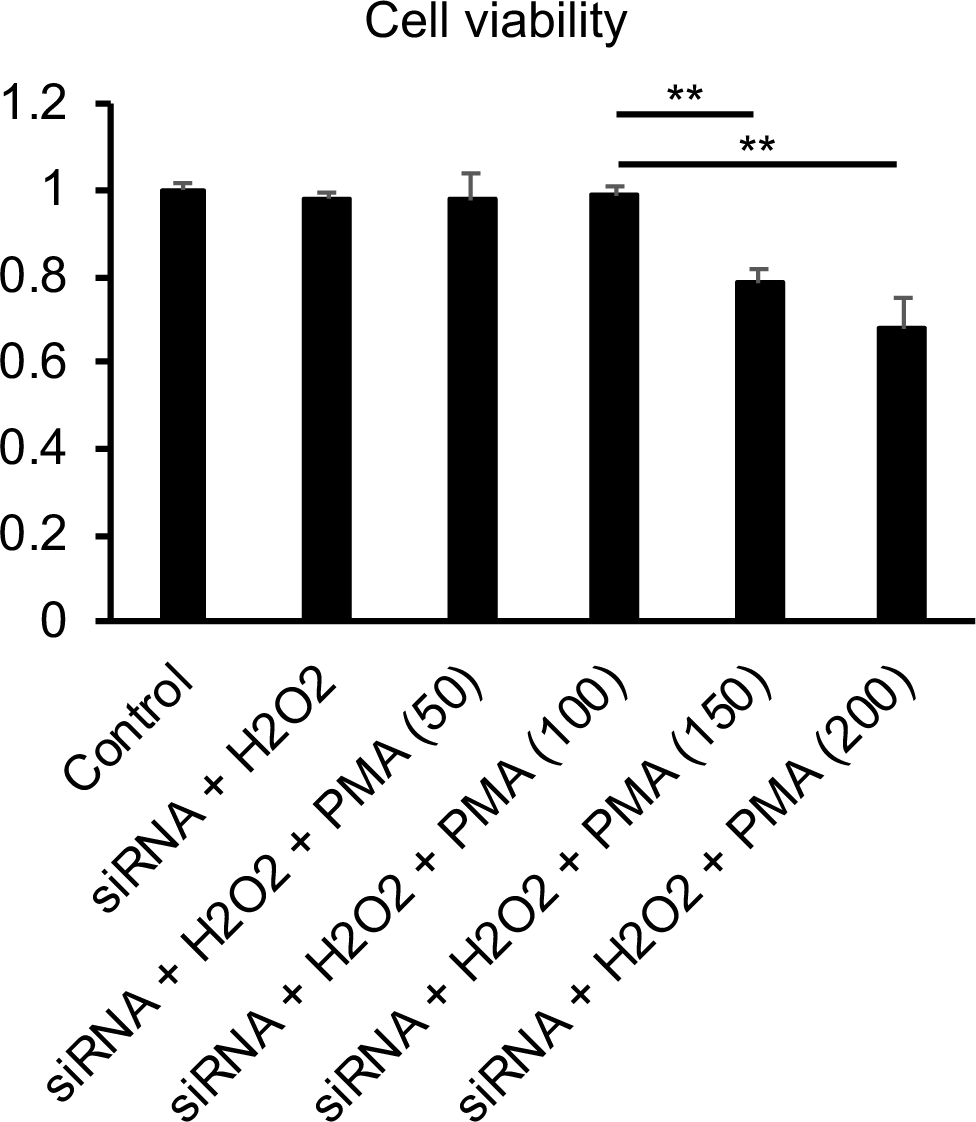

**Supplemental Figure 2.**
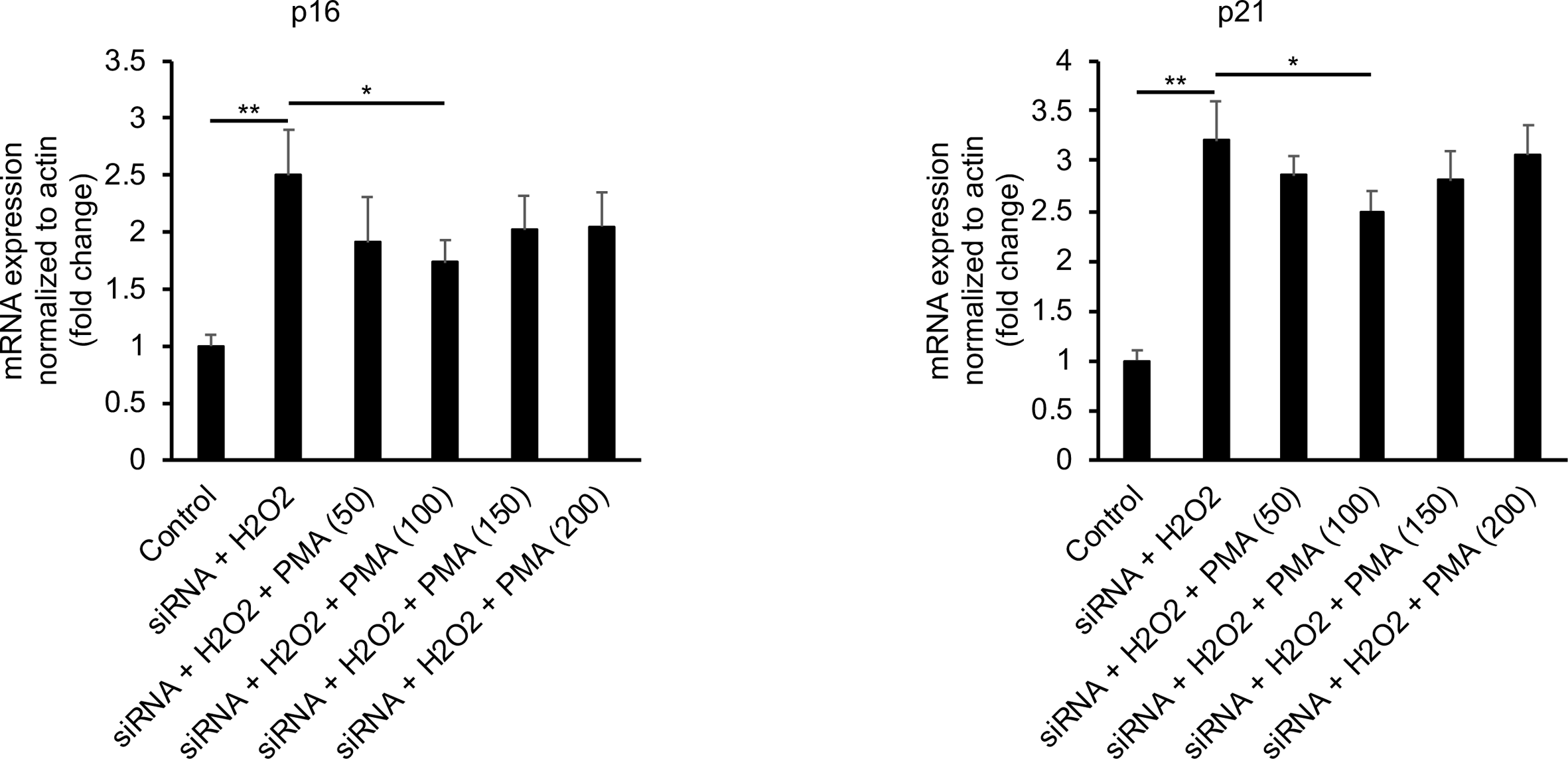

## References

[1] R.S. Apte, Age-Related Macular Degeneration, N Engl J Med 385(6) (2021) 539–547.

[2] P. Mitchell, G. Liew, B. Gopinath, T.Y. Wong, Age-related macular degeneration, Lancet 392(10153) (2018) 1147–1159.

[3] L.G. Fritsche, W. Igl, J.N. Bailey, F. Grassmann, S. Sengupta, J.L. Bragg-Gresham, K.P. Burdon, S.J. Hebbring, C. Wen, M. Gorski, I.K. Kim, D. Cho, D. Zack, E. Souied, H.P. Scholl, E. Bala, K.E. Lee, D.J. Hunter, R.J. Sardell, P. Mitchell, J.E. Merriam, V. Cipriani, J.D. Hoffman, T. Schick, Y.T. Lechanteur, R.H. Guymer, M.P. Johnson, Y. Jiang, C.M. Stanton, G.H. Buitendijk, X. Zhan, A.M. Kwong, A. Boleda, M. Brooks, L. Gieser, R. Ratnapriya, K.E. Branham, J.R. Foerster, J.R. Heckenlively, M.I. Othman, B.J. Vote, H.H. Liang, E. Souzeau, I.L. McAllister, T. Isaacs, J. Hall, S. Lake, D.A. Mackey, I.J. Constable, J.E. Craig, T.E. Kitchner, Z. Yang, Z. Su, H. Luo, D. Chen, H. Ouyang, K. Flagg, D. Lin, G. Mao, H. Ferreyra, K. Stark, C.N. von Strachwitz, A. Wolf, C. Brandl, G. Rudolph, M. Olden, M.A. Morrison, D.J. Morgan, M. Schu, J. Ahn, G. Silvestri, E.E. Tsironi, K.H. Park, L.A. Farrer, A. Orlin, A. Brucker, M. Li, C.A. Curcio, S. Mohand-Said, J.A. Sahel, I. Audo, M. Benchaboune, A.J. Cree, C.A. Rennie, S.V. Goverdhan, M. Grunin, S. Hagbi-Levi, P. Campochiaro, N. Katsanis, F.G. Holz, F. Blond, H. Blanche, J.F. Deleuze, R.P. Igo, Jr., B. Truitt, N.S. Peachey, S.M. Meuer, C.E. Myers, E.L. Moore, R. Klein, M.A. Hauser, E.A. Postel, M.D. Courtenay, S.G. Schwartz, J.L. Kovach, W.K. Scott, G. Liew, A.G. Tan, B. Gopinath, J.C. Merriam, R.T. Smith, J.C. Khan, H. Shahid, A.T. Moore, J.A. McGrath, R. Laux, M.A. Brantley, Jr., A. Agarwal, L. Ersoy, A. Caramoy, T. Langmann, N.T. Saksens, E.K. de Jong, C.B. Hoyng, M.S. Cain, A.J. Richardson, T.M. Martin, J. Blangero, D.E. Weeks, B. Dhillon, C.M. van Duijn, K.F. Doheny, J. Romm, C.C. Klaver, C. Hayward, M.B. Gorin, M.L. Klein, P.N. Baird, A.I. den Hollander, S. Fauser, J.R. Yates, R. Allikmets, J.J. Wang, D.A. Schaumberg, B.E. Klein, S.A. Hagstrom, I. Chowers, A.J. Lotery, T. Leveillard, K. Zhang, M.H. Brilliant, A.W. Hewitt, A. Swaroop, E.Y. Chew, M.A. Pericak-Vance, M. DeAngelis, D. Stambolian, J.L. Haines, S.K. Iyengar, B.H. Weber, G.R. Abecasis, I.M. Heid, A large genome-wide association study of age-related macular degeneration highlights contributions of rare and common variants, Nat Genet 48(2) (2016) 134–43.

[4] S. Arakawa, A. Takahashi, K. Ashikawa, N. Hosono, T. Aoi, M. Yasuda, Y. Oshima, S. Yoshida, H. Enaida, T. Tsuchihashi, K. Mori, S. Honda, A. Negi, A. Arakawa, K. Kadonosono, Y. Kiyohara, N. Kamatani, Y. Nakamura, T. Ishibashi, M. Kubo, Genome-wide association study identifies two susceptibility loci for exudative age-related macular degeneration in the Japanese population, Nat Genet 43(10) (2011) 1001–4.

[5] M. Akiyama, M. Miyake, Y. Momozawa, S. Arakawa, M. Maruyama-Inoue, M. Endo, Y. Iwasaki, K. Ishigaki, N. Matoba, Y. Okada, M. Yasuda, Y. Oshima, S. Yoshida, S.Y. Nakao, K. Morino, Y. Mori, A. Kido, A. Kato, T. Yasukawa, R. Obata, Y. Nagai, K. Takahashi, K. Fujisawa, A. Miki, M. Nakamura, S. Honda, H. Ushida, T. Yasuma, K.M. Nishiguchi, R. Mori, K. Tanaka, Y. Wakatsuki, K. Yamashiro, K. Kadonosono, C. Terao, T. Ishibashi, A. Tsujikawa, K.H. Sonoda, M. Kubo, Y. Kamatani, Genome-Wide Association Study of Age-Related Macular Degeneration Reveals 2 New Loci Implying Shared Genetic Components with Central Serous Chorioretinopathy, Ophthalmology 130(4) (2023) 361–372.

[6] K. Mori, K. Ishikawa, Y. Fukuda, R. Ji, I. Wada, Y. Kubo, M. Akiyama, S. Notomi, Y. Murakami, S. Nakao, S. Arakawa, S. Shiose, T. Hisatomi, S. Yoshida, R. Kannan, K.H. Sonoda, TNFRSF10A downregulation induces retinal pigment epithelium degeneration during the pathogenesis of age-related macular degeneration and central serous chorioretinopathy, Hum Mol Genet 31(13) (2022) 2194–2206.

[7] J.B. Chae, H. Jang, C. Son, C.W. Park, H. Choi, S. Jin, H.Y. Lee, H. Lee, J.H. Ryu, N. Kim, C. Kim, H. Chung, Targeting senescent retinal pigment epithelial cells facilitates retinal regeneration in mouse models of age-related macular degeneration, Geroscience 43(6) (2021) 2809–2833.

[8] P.G. Sreekumar, S.T. Reddy, D.R. Hinton, R. Kannan, Mechanisms of RPE senescence and potential role of alphaB crystallin peptide as a senolytic agent in experimental AMD, Exp Eye Res 215 (2022) 108918.

[9] C. Cavallotti, M. Artico, N. Pescosolido, F.M. Leali, J. Feher, Age-related changes in the human retina, Can J Ophthalmol 39(1) (2004) 61–8.

[10] M.A. Samuel, Y. Zhang, M. Meister, J.R. Sanes, Age-related alterations in neurons of the mouse retina, J Neurosci 31(44) (2011) 16033–44.

[11] P.G. Sreekumar, D.R. Hinton, R. Kannan, The Emerging Role of Senescence in Ocular Disease, Oxid Med Cell Longev 2020 (2020) 2583601.

[12] H. Gao, J.G. Hollyfield, Aging of the human retina. Differential loss of neurons and retinal pigment epithelial cells, Invest Ophthalmol Vis Sci 33(1) (1992) 1–17.

[13] C.K. Dorey, G. Wu, D. Ebenstein, A. Garsd, J.J. Weiter, Cell loss in the aging retina. Relationship to lipofuscin accumulation and macular degeneration, Invest Ophthalmol Vis Sci 30(8) (1989) 1691–9.

[14] E. Chaum, C.S. Winborn, S. Bhattacharya, Genomic regulation of senescence and innate immunity signaling in the retinal pigment epithelium, Mamm Genome 26(5-6) (2015) 210–21.

[15] H. Matsunaga, J.T. Handa, A. Aotaki-Keen, S.W. Sherwood, M.D. West, L.M. Hjelmeland, Beta-galactosidase histochemistry and telomere loss in senescent retinal pigment epithelial cells, Invest Ophthalmol Vis Sci 40(1) (1999) 197–202.

[16] K. Mishima, J.T. Handa, A. Aotaki-Keen, G.A. Lutty, L.S. Morse, L.M. Hjelmeland, Senescence-associated beta-galactosidase histochemistry for the primate eye, Invest Ophthalmol Vis Sci 40(7) (1999) 1590–3.

[17] J.R. Sparrow, M. Boulton, RPE lipofuscin and its role in retinal pathobiology, Exp Eye Res 80(5) (2005) 595–606.

[18] P.G. Sreekumar, K. Ishikawa, C. Spee, H.H. Mehta, J. Wan, K. Yen, P. Cohen, R. Kannan, D.R. Hinton, The Mitochondrial-Derived Peptide Humanin Protects RPE Cells From Oxidative Stress, Senescence, and Mitochondrial Dysfunction, Invest Ophthalmol Vis Sci 57(3) (2016) 1238–53.

[19] K. Ishikawa, P.G. Sreekumar, C. Spee, H. Nazari, D. Zhu, R. Kannan, D.R. Hinton, alphaB-Crystallin Regulates Subretinal Fibrosis by Modulation of Epithelial-Mesenchymal Transition, Am J Pathol 186(4) (2016) 859–73.

[20] Z. Chen, C. Zhang, X. Song, X. Cui, J. Liu, N.C. Ford, S. He, G. Zhu, X. Dong, M. Hanani, Y. Guan, BzATP Activates Satellite Glial Cells and Increases the Excitability of Dorsal Root Ganglia Neurons In Vivo, Cells 11(15) (2022).

[21] K. Elgass, J. Pakay, M.T. Ryan, C.S. Palmer, Recent advances into the understanding of mitochondrial fission, Biochim Biophys Acta 1833(1) (2013) 150–61.

[22] C. Hu, Y. Huang, L. Li, Drp1-Dependent Mitochondrial Fission Plays Critical Roles in Physiological and Pathological Progresses in Mammals, Int J Mol Sci 18(1) (2017).

[23] A.P. Gureev, E.A. Shaforostova, V.N. Popov, Regulation of Mitochondrial Biogenesis as a Way for Active Longevity: Interaction Between the Nrf2 and PGC-1alpha Signaling Pathways, Front Genet 10 (2019) 435.

[24] C. Kukat, N.G. Larsson, mtDNA makes a U-turn for the mitochondrial nucleoid, Trends Cell Biol 23(9) (2013) 457–63.

[25] J.R. Dorr, Y. Yu, M. Milanovic, G. Beuster, C. Zasada, J.H. Dabritz, J. Lisec, D. Lenze, A. Gerhardt, K. Schleicher, S. Kratzat, B. Purfurst, S. Walenta, W. Mueller-Klieser, M. Graler, M. Hummel, U. Keller, A.K. Buck, B. Dorken, L. Willmitzer, M. Reimann, S. Kempa, S. Lee, C.A. Schmitt, Synthetic lethal metabolic targeting of cellular senescence in cancer therapy, Nature 501(7467) (2013) 421–5.

[26] T. Kamogashira, K. Hayashi, C. Fujimoto, S. Iwasaki, T. Yamasoba, Functionally and morphologically damaged mitochondria observed in auditory cells under senescence-inducing stress, NPJ Aging Mech Dis 3 (2017) 2.

[27] S.J. Kim, H.H. Mehta, J. Wan, C. Kuehnemann, J. Chen, J.F. Hu, A.R. Hoffman, P. Cohen, Mitochondrial peptides modulate mitochondrial function during cellular senescence, Aging (Albany NY) 10(6) (2018) 1239–1256.

[28] J. Martinez, D. Tarallo, L. Martinez-Palma, S. Victoria, M. Bresque, S. Rodriguez-Bottero, I. Marmisolle, C. Escande, P. Cassina, G. Casanova, M. Bollati-Fogolin, C. Agorio, M. Moreno, C. Quijano, Mitofusins modulate the increase in mitochondrial length, bioenergetics and secretory phenotype in therapy-induced senescent melanoma cells, Biochem J 476(17) (2019) 2463–2486.

[29] J.M.T. Hyttinen, J. Viiri, K. Kaarniranta, J. Blasiak, Mitochondrial quality control in AMD: does mitophagy play a pivotal role?, Cell Mol Life Sci 75(16) (2018) 2991–3008.

[30] N. Basisty, A. Kale, O.H. Jeon, C. Kuehnemann, T. Payne, C. Rao, A. Holtz, S. Shah, V. Sharma, L. Ferrucci, J. Campisi, B. Schilling, A proteomic atlas of senescence-associated secretomes for aging biomarker development, PLoS Biol 18(1) (2020) e3000599.

[31] J.P. Coppe, P.Y. Desprez, A. Krtolica, J. Campisi, The senescence-associated secretory phenotype: the dark side of tumor suppression, Annu Rev Pathol 5 (2010) 99–118.

[32] T. Tchkonia, Y. Zhu, J. van Deursen, J. Campisi, J.L. Kirkland, Cellular senescence and the senescent secretory phenotype: therapeutic opportunities, J Clin Invest 123(3) (2013) 966–72.

[33] K.S. Lee, S. Lin, D.A. Copland, A.D. Dick, J. Liu, Cellular senescence in the aging retina and developments of senotherapies for age-related macular degeneration, J Neuroinflammation 18(1) (2021) 32.

[34] J.D. Mott, Z. Werb, Regulation of matrix biology by matrix metalloproteinases, Curr Opin Cell Biol 16(5) (2004) 558–64.

[35] J.T. Handa, How does the macula protect itself from oxidative stress?, Mol Aspects Med 33(4) (2012) 418–35.

[36] S.G. Jarrett, M.E. Boulton, Consequences of oxidative stress in age-related macular degeneration, Mol Aspects Med 33(4) (2012) 399–417.

[37] L.G. Fritsche, W. Chen, M. Schu, B.L. Yaspan, Y. Yu, G. Thorleifsson, D.J. Zack, S. Arakawa, V. Cipriani, S. Ripke, R.P. Igo, Jr., G.H. Buitendijk, X. Sim, D.E. Weeks, R.H. Guymer, J.E. Merriam, P.J. Francis, G. Hannum, A. Agarwal, A.M. Armbrecht, I. Audo, T. Aung, G.R. Barile, M. Benchaboune, A.C. Bird, P.N. Bishop, K.E. Branham, M. Brooks, A.J. Brucker, W.H. Cade, M.S. Cain, P.A. Campochiaro, C.C. Chan, C.Y. Cheng, E.Y. Chew, K.A. Chin, I. Chowers, D.G. Clayton, R. Cojocaru, Y.P. Conley, B.K. Cornes, M.J. Daly, B. Dhillon, A.O. Edwards, E. Evangelou, J. Fagerness, H.A. Ferreyra, J.S. Friedman, A. Geirsdottir, R.J. George, C. Gieger, N. Gupta, S.A. Hagstrom, S.P. Harding, C. Haritoglou, J.R. Heckenlively, F.G. Holz, G. Hughes, J.P. Ioannidis, T. Ishibashi, P. Joseph, G. Jun, Y. Kamatani, N. Katsanis, N.K. C J.C. Khan, I.K. Kim, Y. Kiyohara, B.E. Klein, R. Klein, J.L. Kovach, I. Kozak, C.J. Lee, K.E. Lee, P. Lichtner, A.J. Lotery, T. Meitinger, P. Mitchell, S. Mohand-Said, A.T. Moore, D.J. Morgan, M.A. Morrison, C.E. Myers, A.C. Naj, Y. Nakamura, Y. Okada, A. Orlin, M.C. Ortube, M.I. Othman, C. Pappas, K.H. Park, G.J. Pauer, N.S. Peachey, O. Poch, R.R. Priya, R. Reynolds, A.J. Richardson, R. Ripp, G. Rudolph, E. Ryu, J.A. Sahel, D.A. Schaumberg, H.P. Scholl, S.G. Schwartz, W.K. Scott, H. Shahid, H. Sigurdsson, G. Silvestri, T.A. Sivakumaran, R.T. Smith, L. Sobrin, E.H. Souied, D.E. Stambolian, H. Stefansson, G.M. Sturgill-Short, A. Takahashi, N. Tosakulwong, B.J. Truitt, E.E. Tsironi, A.G. Uitterlinden, C.M. van Duijn, L. Vijaya, J.R. Vingerling, E.N. Vithana, A.R. Webster, H.E. Wichmann, T.W. Winkler, T.Y. Wong, A.F. Wright, D. Zelenika, M. Zhang, L. Zhao, K. Zhang, M.L. Klein, G.S. Hageman, G.M. Lathrop, K. Stefansson, R. Allikmets, P.N. Baird, M.B. Gorin, J.J. Wang, C.C. Klaver, J.M. Seddon, M.A. Pericak-Vance, S.K. Iyengar, J.R. Yates, A. Swaroop, B.H. Weber, M. Kubo, M.M. Deangelis, T. Leveillard, U. Thorsteinsdottir, J.L. Haines, L.A. Farrer, I.M. Heid, G.R. Abecasis, A.M.D.G. Consortium, Seven new loci associated with age-related macular degeneration, Nat Genet 45(4) (2013) 433–9, 439e1-2.

[38] K. Kaarniranta, J. Kajdanek, J. Morawiec, E. Pawlowska, J. Blasiak, PGC-1alpha Protects RPE Cells of the Aging Retina against Oxidative Stress-Induced Degeneration through the Regulation of Senescence and Mitochondrial Quality Control. The Significance for AMD Pathogenesis, Int J Mol Sci 19(8) (2018).

[39] K. Kaarniranta, E. Pawlowska, J. Szczepanska, A. Jablkowska, J. Blasiak, Role of Mitochondrial DNA Damage in ROS-Mediated Pathogenesis of Age-Related Macular Degeneration (AMD), Int J Mol Sci 20(10) (2019).

[40] B.N. Ames, M.K. Shigenaga, Oxidants are a major contributor to aging, Ann N Y Acad Sci 663 (1992) 85–96.

[41] N.T. Moldogazieva, I.M. Mokhosoev, T.I. Mel’nikova, Y.B. Porozov, A.A. Terentiev, Oxidative Stress and Advanced Lipoxidation and Glycation End Products (ALEs and AGEs) in Aging and Age-Related Diseases, Oxid Med Cell Longev 2019 (2019) 3085756.

[42] D.A. Ferrington, R.J. Kapphahn, M.M. Leary, S.R. Atilano, M.R. Terluk, P. Karunadharma, G.K. Chen, R. Ratnapriya, A. Swaroop, S.R. Montezuma, M.C. Kenney, Increased retinal mtDNA damage in the CFH variant associated with age-related macular degeneration, Exp Eye Res 145 (2016) 269–277.

[43] J. Blasiak, S. Glowacki, A. Kauppinen, K. Kaarniranta, Mitochondrial and nuclear DNA damage and repair in age-related macular degeneration, Int J Mol Sci 14(2) (2013) 2996–3010.

[44] H. Rottenberg, J.B. Hoek, The path from mitochondrial ROS to aging runs through the mitochondrial permeability transition pore, Aging Cell 16(5) (2017) 943–955.

[45] M. Ji Cho, S.J. Yoon, W. Kim, J. Park, J. Lee, J.G. Park, Y.L. Cho, J. Hun Kim, H. Jang, Y.J. Park, S.H. Lee, J.K. Min, Oxidative stress-mediated TXNIP loss causes RPE dysfunction, Exp Mol Med 51(10) (2019) 1–13.

[46] M. Kitada, D. Koya, Autophagy in metabolic disease and ageing, Nat Rev Endocrinol 17(11) (2021) 647–661.

[47] K. Kaarniranta, H. Uusitalo, J. Blasiak, S. Felszeghy, R. Kannan, A. Kauppinen, A. Salminen, D. Sinha, D. Ferrington, Mechanisms of mitochondrial dysfunction and their impact on age-related macular degeneration, Prog Retin Eye Res 79 (2020) 100858.

[48] Z. Lu, D. Liu, A. Hornia, W. Devonish, M. Pagano, D.A. Foster, Activation of protein kinase C triggers its ubiquitination and degradation, Mol Cell Biol 18(2) (1998) 839–45.

[49] C. Michaloglou, L.C. Vredeveld, M.S. Soengas, C. Denoyelle, T. Kuilman, C.M. van der Horst, D.M. Majoor, J.W. Shay, W.J. Mooi, D.S. Peeper, BRAFE600-associated senescence-like cell cycle arrest of human naevi, Nature 436(7051) (2005) 720–4.

[50] Y. Mori, A.K. Ajay, J.H. Chang, S. Mou, H. Zhao, S. Kishi, J. Li, C.R. Brooks, S. Xiao, H.M. Woo, V.S. Sabbisetti, S.C. Palmer, P. Galichon, L. Li, J.M. Henderson, V.K. Kuchroo, J. Hawkins, T. Ichimura, J.V. Bonventre, KIM-1 mediates fatty acid uptake by renal tubular cells to promote progressive diabetic kidney disease, Cell Metab 33(5) (2021) 1042–1061 e7.

