## Supplementary material for "Characterization and contribution of RPE senescence to Age-related macular degeneration in *Tnfrsf10* knock out mice": Table for primer

Table 1. Primer Sequences Used for Real-Time RT-PCR

| Target molecule | Forward primer sequence | Reverse primer sequence |
|---|---|---|
| p21 | 5`-AAG TCA GTT CCT TGT GGA GC-3` | 5`GCC ATT AGC GCA TCA CAG TC-3` |
| p16 | GTG GGT TTG TAG AAG CAG GCA | ATC CCC AGG CAT CTT TTG CAC |
| IFN-γ | 5`-TGA CCA GAG CAT CCA AAA GA-3` | 5`-CTC TTC GAC CTC GAA ACA GC-3` |
| IL-1β | 5`-GGC CCT AAA CAG ATG AAG TGCT-3` | 5`-TGC CGC CAT CCA GAG G-3` |
| IL-6 | 5`-AAT TCG GTA CAT CCT CGA CGG CGG-3` | 5`-GGT TGT TTT CTG CCA GTG CC-3` |
| IL-10 | 5`-GGT TGC CAA GCC TTG TCT GA-3` | 5`-AGG GAG TTC ACA TGC GCC T-3` |
| IL-11 | 5`-GTG GCC AGA TAC AGC TGT CGC-3` | 5`-GGT AGG ACA GTA GGT CCG CTC-3` |
| MMP-1 | 5`-GCT AAC CTT TGA TGC TAT AAC TAC GA-3` | 5`-TTT GTG CGC ATG TAG AAT CTG-3` |
| MMP-3 | 5`-CCA GGT GTG GAG TTC CTG AT-3` | 5`-CAT CTT TTG GCA AAT CTG GTG-3` |
| Tubulin | 5`-CTT CGT CTC CGC CAT CAG-3` | 5`-CGT GTT CCA GGC AGT AGA GC-3` |
| Actin | 5`-CAC CAT TGG CAA TGA GCG GTT C-3` | 5`-AGG TCT TTG CGG ATG TCC ACG T-3` |

Table 2. List of Antibodies Used for Western Blotting

| Target molecule | Source | Dilution | Product catalogue |
|---|---|---|---|
| p16$^{INK4a}$ | Rabbit monoclonal | 1: 1000 | Abcam (ab108349) |
| p21 | Mouse | 1: 1000 | Santa Cruz Biotech (sc-6246) |
| Fis1 | Mouse | 1: 1000 | Santa Cruz Biotech (sc-376447) |
| DRP1 | Mouse | 1: 1000 | Santa Cruz Biotech (sc-271583) |
| PGC1-α | Mouse | 1: 1000 | Santa Cruz Biotech (sc-518038) |
| mtTFA | Mouse | 1: 1000 | Santa Cruz Biotech (sc-166965) |
| GAPDH (MAB374) | Mouse | 1: 1000 | EMD Millipore (MAB374) |
